# Reinforced Hippocampus-Accumbens pathway marks a shift towards cognitive decline in aging

**DOI:** 10.64898/2026.01.19.700256

**Authors:** Aurore Ribera, Brenda Fernandes-Dias, Larissa Yugay, Emma Deneuville, Manuel Dias-Silva, Marion Violain, Thibault Bouet, Ingrid Bethus, Luisa V. Lopes, Peter Vanhoutte, Stephen Ramanoël, Jacques Barik, Paula A. Pousinha

## Abstract

Aging triggers early functional alterations in brain circuits that precede cognitive impairment, yet the underlying mechanisms remain unclear. Spatial navigation and memory, among the earliest affected, depend on hippocampus and downstream targets such as the nucleus accumbens (NAc), which integrates spatial and motivational information to guide goal-directed behaviors. Here we identify age-related, sex-dependent changes of the dorsal hippocampus–NAc (dHPC→NAc) pathway. Combining optogenetics with electrophysiology, chemogenetics, and behavioral analyses in mice, we show that aging shifts excitation–inhibition balance in dorsal CA1 toward pyramidal neuron hyperexcitability, strengthening hippocampal outputs. This selectively enhances synaptic drive onto D1 receptor-expressing medium spiny neurons and involves a previously unrecognized long-range parvalbumin-expressing glutamatergic projection. Reducing dHPC→NAc excitability improves memory in aged mice. In humans, fMRI reveals heightened posterior hippocampal activity and stronger pHPC–NAc coupling during navigation. Together, we uncover a targetable pathway acting in an intervention-sensitive window of aging to correct negative cognitive trajectories.

## Introduction

Aging involves widespread changes in brain structures and functions that emerge well before clinical symptoms become apparent^1^. However, cognitive aging follows a heterogeneous pattern; while some individuals maintain preserved cognitive abilities, others experience significant decline. Evidence from neuroimaging and neuropathological studies suggests that brain changes start to appear around 50 - 60 years of age, identifying this as a critical window for early intervention^2–4^. However, circuit-level studies of brain physiology at this critical age remain scarce. Among the earliest cognitive processes to decline are spatial navigation, including in reward seeking tasks, and memory^3,5,6^, with sex specificities^7^. These functions that depend heavily on the hippocampus (HPC)^8–14^. The hippocampus (HPC), composed of subregions^15–17^ with distinct connectivity and cellular profiles, is one of the most extensively studied regions in healthy brain aging^18–20^ due to its pronounced vulnerability to age-related structural and functional deterioration. Human studies consistently show hippocampal volume decrease upon aging^5,21–23^, with some reporting sex differences^24,25^. Within the HPC, CA1 pyramidal neurons, which constitute its primary output^26,27^, play a critical role in cognitive function but are especially vulnerable to age-related alterations^28–30^. Aging is associated with heightened excitability in the HPC^31,32^, suggesting increased hippocampal output to connected regions, though the impact of these alterations on HPC-target regions remains poorly understood. One such target, the nucleus accumbens (NAc), exhibit decreased volume during aging^33–36^. The NAc, is a key brain region that integrates diverse information from several brain areas to effectively mediate goal-directed behaviors^37^. Yet, patients with volumetric decrease in both structures display worse cognitive scores and are more susceptible to develop neurodegenerative disorders, such as Alzheimer’s disease^38^. The HPC projects to the NAc^39,40^, forming a circuit that integrates spatial and motivational information, suggesting that alterations in this pathway could contribute to age-related deficits in goal-directed behaviors. Furthermore, since these brain structures are both vulnerable to aging at the critical phase of early aging, age-related changes in this neuronal pathway may constitute an early biomarker of cognitive decline.

By combining anatomical tracing, optogenetics coupled to electrophysiology, chemogenetics and behavioral analyses, we show that aging shifts the excitation-inhibition balance in dorsal CA1 toward increased excitation, producing hippocampal hyperactivity in this major output region. Within the dHPC^→NAc^pathway, we identify a previously unrecognized long-range fast-spiking population co-expressing parvalbumin and glutamatergic markers (PV-Glut). Circuit interrogation further reveals that monosynaptic dHPC inputs differentially modulate NAc cell populations. It enhances synaptic drive onto dopamine D1 receptor–expressing medium spiny neurons (D1R-MSN) while exerting opposing effects on D2R-MSN. Notably, stronger dHPC^→D1R-MSN^transmission is negatively associated with reward-driven spatial memory performance. Selective chemogenetic dampening of dHPC^→NAc^projecting neurons hyperactivity improves reward-based spatial memory, specifically in aged mice. In line with these findings, human neuroimaging in healthy older adults reveals increased hippocampal activity and enhanced hippocampus-NAc coupling during navigation. Together, our data identify a circuit mechanism by which age-related hippocampal hyperactivity impacts the NAc function and associated reward driven spatial memory, providing a framework for circuit-informed interventions to preserve cognitive health span.

## Results

### CA1 output neurons show increased synaptic excitability in aging

Aligned with previous studies showing that aging is associated with heightened excitability in the HPC^31,32^, we investigated the excitatory/inhibitory (E/I) ratio of the output CA1 pyramidal neurons in hippocampal slices from aged mice. We observed a pronounced shift in the E/I balance towards increased excitation in neurons from aged animals (Fig. 1a-d). We thus investigated whether this E/I balance disruption may affect dHPC cells projecting to the NAc (dHPC^→NAc^). In order to perform a multivariable analysis per animal, the experimental design followed the chronology shown in Fig. 1e. To unambiguously identify this neuronal pathway, we employed an intersectional viral tracing with retrograde CAV2-Cre injection in the NAc combined with Cre-dependent expression of fluorescent marker mCherry (AAV-DIO-mCherry) in the dHPC (Fig. 1f(i)). This specific labelling allowed the analyses of the anatomical distribution and electrophysiological profiling of dHPC^→NAc^neurons. We found sparse labelling thus suggesting that this projecting neuronal population constitutes a small subset within the HPC CA1 subregion (Fig. 1f(ii)). Electrophysiological recordings of their intrinsic properties and excitability showed no significant age-related differences (Fig. 1g,h; Extended data Fig. 1). Upon depolarizing current injections, we unveiled two types of firing patterns among CA1 neurons: one group exhibited responses consistent with typical pyramidal cell profiles (Fig. 1g, regular spiking) while a second population showed high-frequency firing approaching ∼150Hz, reminiscent of parvalbumin (PV)-expressing fast spiking neurons (Fig. 1h, fast spiking). This functional evidence was complemented by immunohistofluorescence analyses that reveal co-expression of PV marker and mCherry (Fig. 1i). These findings suggest a heterogeneous dHPC^→NAc^neuronal population composed of pyramidal cells and long-range PV-expressing neurons.

**Figure 1:**
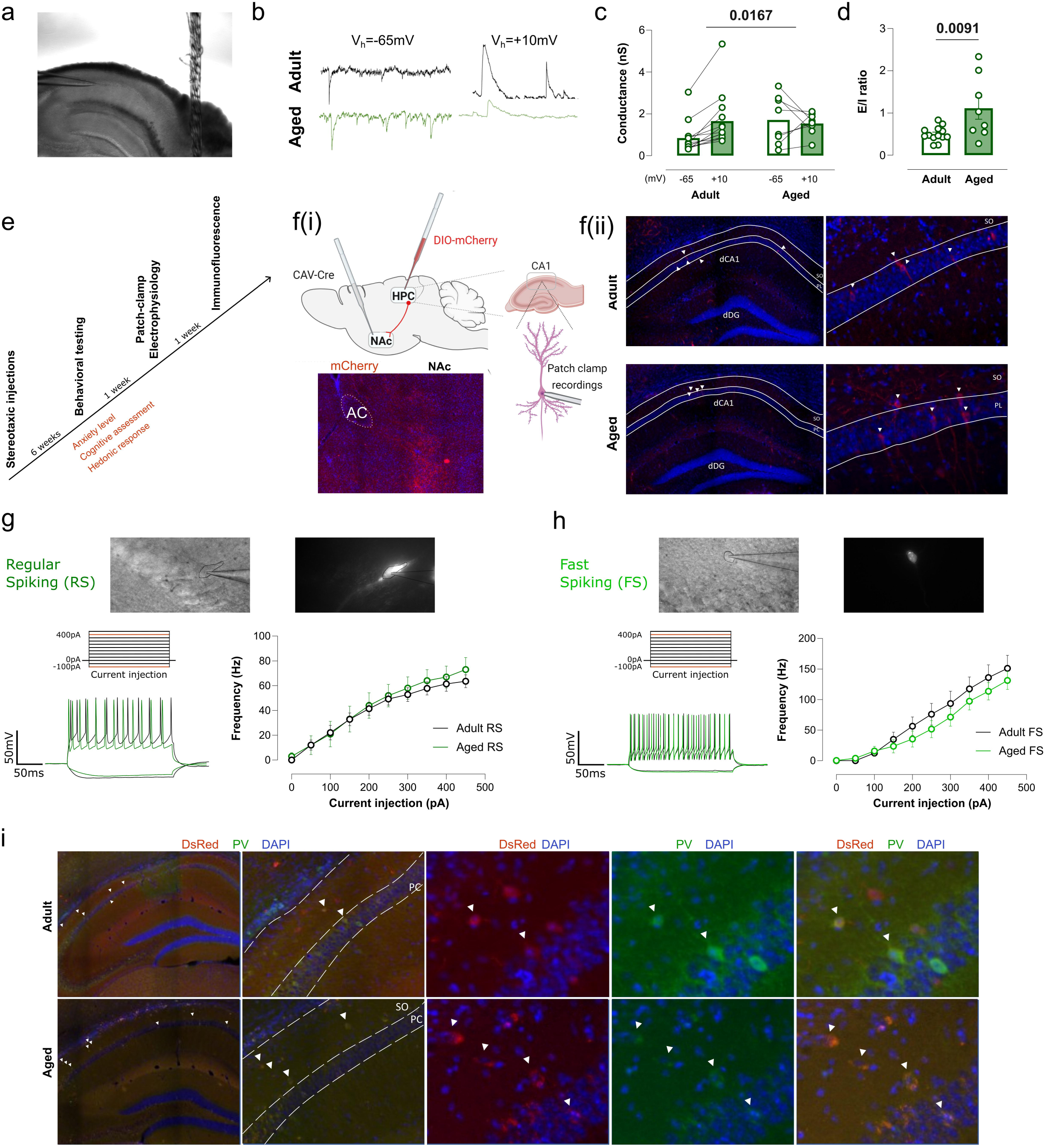
CA1 output neurons show increased synaptic excitability in aging. (a) **Representative image of hippocampal slice with the pipette on CA1 region, 100x magnification.** **(b)** Illustrative traces of recorded spontaneous currents. **(c)** Synaptic conductances of CA1 pyramidal cells were derived from currents recorded at holding potentials of −65 mV for excitatory and +10 mV for inhibitory inputs. Excitatory conductance is increased in aging, and not followed by inhibitory responses. Results are expressed as the mean ± SEM (Two-way ANOVA, interaction: p = 0.0167, F(1, 19) = 6.892, adult N = 4, n = 13, aged N = 4, n = 8). **(d)** The balance excitation/inhibition (E/I ratio) is increased in aging towards excitation. Results are expressed as the mean ± SEM (Passed normality test. Unpaired t-test, p = 0.0091, adult N = 4, n = 13, aged N = 4, n = 8). **(e)** Experimental design. **(f(i))** Schematic representation of stereotaxic injections: intersectional viral tracing with retrograde CAV2-Cre injection in the NAc combined with Cre-dependent expression of AAV-DIO-mCherry in the dHPC. At the bottom, illustrative image of the fibres of the dHPC projecting-cells in the NAc. **(f(ii))** Representative images of the labelling of the dHPC cells projecting to the NAc at two amplifications. dDG = dorsal dentate gyrus, SO = stratum oriens, PL = pyramidal layer, AC = anterior commissura, **(g)** Representative images of the CA1 regular-spiking (RS, recorded cells with a recording pipette, without and with fluorescence (600x magnification). Protocol used to assess intrinsic excitability properties of the cells. Comparison of representative whole-cell patch-clamp recordings of excitability profile from adult (3-4 months), and aged (18–20 months) C57BL/6 wild-type mice, illustrating no differences, confirmed in the graph showing the excitability profile. Results are expressed as the mean ± SEM (Two-way ANOVA with repeated measures, age: p = 0.7010, F(1, 10) = 0.1561, adult N= 5, n = 7, aged N= 4, n = 5). **(h)** Same as (g) but for dHPC fast-spiking (FS) cells. Results are expressed as the mean ± SEM (Two-way ANOVA with repeated measures, age: p = 0.4129, F(1, 19) = 0.7008, adult N = 7, n = 10, aged N = 7, n = 11). **(i)** Representative immunocytochemistry analysis of Hippocampal CA1 neurons projecting to the NAc in hippocampal slices from both adult and aged mice (N= 8 per group). Nuclei were stained with DAPI (blue); parvalbumin (PV) expression is shown in green; and DsRed- enhanced mCherry signal (red) reflects retrograde labelling from hM4D-DIO-mCherry virus injected into the NAc. Lower panels show magnified views of the boxed regions in the pyramidal layer and Stratum Oriens of the dHPC. In all panels: significant p-values are represented in bold, N = number of animals, n = number of cells.

### Aging increases spontaneous excitatory synaptic inputs onto D1R-MSN and D2R-MSN

In order to capture age-related effects across principal cell types of the NAc, we evaluated the effects of aging on spontaneous inputs to different cell populations of this brain structure. The NAc is a predominantly GABAergic region composed largely of medium spiny neurons (MSN, 95%) and a smaller population of parvalbumin (PV-expressing, 3%) interneurons^41–43^. We identified the two main NAc populations by selectively recording fluorescently-tagged MSN either D1R or D2R through the use of well-validated viral tools^44^ (Fig. 2a,i). Spontaneous activity of these different cell populations was assessed within the NAc region innervated by the dHPC terminals (Fig. 1f(i), 2b,j). We observed an increased frequency of spontaneous excitatory post-synaptic currents (Fig. 2e, Extended data Fig. 2b), with no change in amplitude (Fig. 2d, Extended data Fig. 2a), thus suggesting heightened excitatory afferent activity onto D1R-MSN. In parallel, spontaneous inhibitory post-synaptic currents showed increased amplitude (Fig. 2g, Extended data Fig. 2c) without frequency changes (Fig. 2h, Extended data Fig. 2d), probably due to post-synaptic homeostatic regulation. Excitatory and inhibitory conductances were similar among groups (Extended data Fig. 2e), maintaining the E/I balance in aging (Extended data Fig. 2f). Similar pattern was observed in D2R-MSN in regard to spontaneous excitatory inputs (Fig. 2k-m, Extended data Fig. 2g,h), but age-related changes in spontaneous inhibitory inputs did not reach statistical significance (Fig. 2n-p, Extended data Fig. 2i-l), thus suggesting different impact of aging on the major NAc cell types. In addition, we also examined the properties of PV-expressing NAc interneurons specifically identified by combining AAV-DIO-mCherry injection in the NAc of PV-Cre mice (Extended data Fig. 3a). PV-expressing neurons appear to be preserved by aging as spontaneous inputs were shown to be similar among groups (Extended data Fig. 3). Together, these results suggest that aging increases spontaneous excitatory synaptic inputs onto D1R-MSN and D2R-MSN, consistent with the increased excitatory drive from the hippocampus.

**Figure 2:**
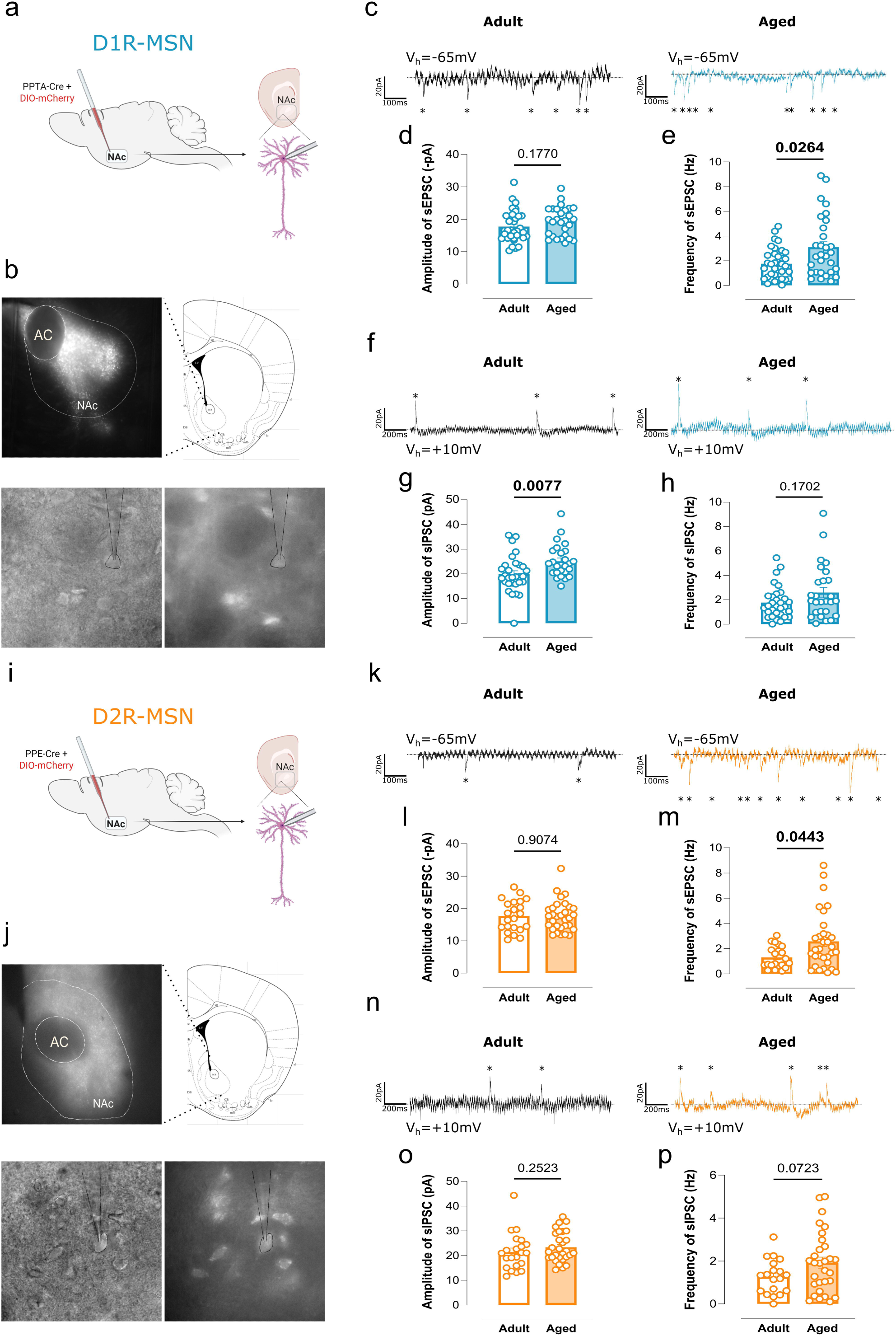
Aging increases spontaneous excitatory synaptic inputs onto D1R-MSN and **D2R-MSN.** **(a)** Schematic representation of stereotaxic injections: viral tracing with PPTA-Cre combined with DIO-mCherry injected in the NAc to label specifically the D1R-MSN population. **(b)** Representative image of the NAc with labelled D1R-MSN population (up: fluorescence in low magnification, 100x, correspondent to the brain atlas coronal slice diagram; down: illustration of a recorded cell with a recording pipette, magnification 600x, without and with fluorescence). AC = anterior commissura. **(c)** Representative traces of spontaneous excitatory post synaptic currents (sEPSC) for adult and aged D1R-MSN. **(d)** Amplitude of sEPSC is similar in both groups. Results are expressed as the mean ± SEM. (Passed normality test. Unpaired t-test, p = 0.1770, adult N = 9, n = 35, aged N = 8, n = 28). **(e)** Frequency of sEPSC is increased in aged D1R-MSN. Results are expressed as the mean ± SEM. (Normality test not passed. Mann-Whitney test, p = 0.0264, adult N = 9, n = 34, aged N = 8, n = 30). **(f)** Representative traces of spontaneous inhibitory post synaptic currents (sIPSC) for adult and aged D1R-MSN. **(g)** Amplitude of sIPSC is increased for aged D1R-MSN. Results are expressed as the mean ± SEM. (Passed normality test. Unpaired t-test, p = 0.0077, adult N = 9, n = 30, aged N = 8, n = 26). **(h)** Frequency of sIPSC is preserved with aging. Results are expressed as the mean ± SEM. (Normality test not passed. Mann-Whitney test, p = 0.1702, adult N = 9, n = 31, aged N = 8, n = 27). **(i)** Schematic representation of stereotaxic injections: viral tracing with PPE-Cre combined with DIO-mCherry injected in the NAc to label specifically the D2R-MSN population. **(j)** Representative image of the NAc with labelled D2R-MSN population (up: fluorescence in low magnification, 100x, correspondent to the brain atlas coronal slice diagram; down: illustration of a recorded cell with a recording pipette, without and with fluorescence, magnification 600x). AC = anterior commissura. **(k)** Representative traces of sEPSC for adult and aged D2R-MSN. **(l)** Amplitude of sEPSC is similar in both groups. Results are expressed as the mean ± SEM. (Normality test not passed. Mann-Whitney test, p = 0.9074, adult N = 7, n = 22, aged N = 10, n = 31). **(m)** Frequency of sEPSC is increased in aged D2R-MSN. Results are expressed as the mean ± SEM. (Normality test not passed. Mann-Whitney test, p = 0.0443, adult N = 7, n = 21, aged N = 10, n = 32). **(n)** Representative traces of sIPSC for adult and aged D2R-MSN. **(o)** Amplitude of sIPSC is maintained in aging for D2R-MSN. Results are expressed as the mean ± SEM. (Normality test not passed. Mann-Whitney test, p = 0.2523, adult N = 7, n = 22, aged N = 10, n = 29). **(p)** Frequency of sIPSC is preserved with aging. Results are expressed as the mean ± SEM. (Passed normality test. Unpaired t-test, p = 0.0723, adult N = 7, n = 19, aged N = 10, n = 30). In all panels: significant p-values are represented in bold, N = number of animals, n = number of cells.

### Aging specifically enhances dHPC^→^**^PV-expressing^ ^neurons^ connectivity**

To investigate synaptic adaptations at dHPC^→NAc^projections, we coupled patch-clamp recordings of identified MSN and PV-expressing neurons, as described above, with optogenetics on NAc brain slices (Fig 3). Channelrhodopsin-2 (AAV-ChR2-mCherry) was injected into the dHPC to enable optical stimulation of hippocampal terminals. Cells showing light-evoked post-synaptic responses were classified as receiving functional dHPC inputs. The three investigated NAc cell populations were shown to be connected to dHPC (Fig. 3c,f,i). Yet, in adults the dHPC^→D1R-MSN^connectivity prevails in comparison to the other cell types, a prevalence that is attenuated in aging in favour to dHPC^→PV-expressing neurons^(Fig. 3c,i). These results show age-related changes within the circuit, with dHPC^→^ ^PV-expressing neurons^ functional connectivity showing the strongest age-related increase, which might result from a homeostatic adaptation to increased excitatory inputs.

**Figure 3:**
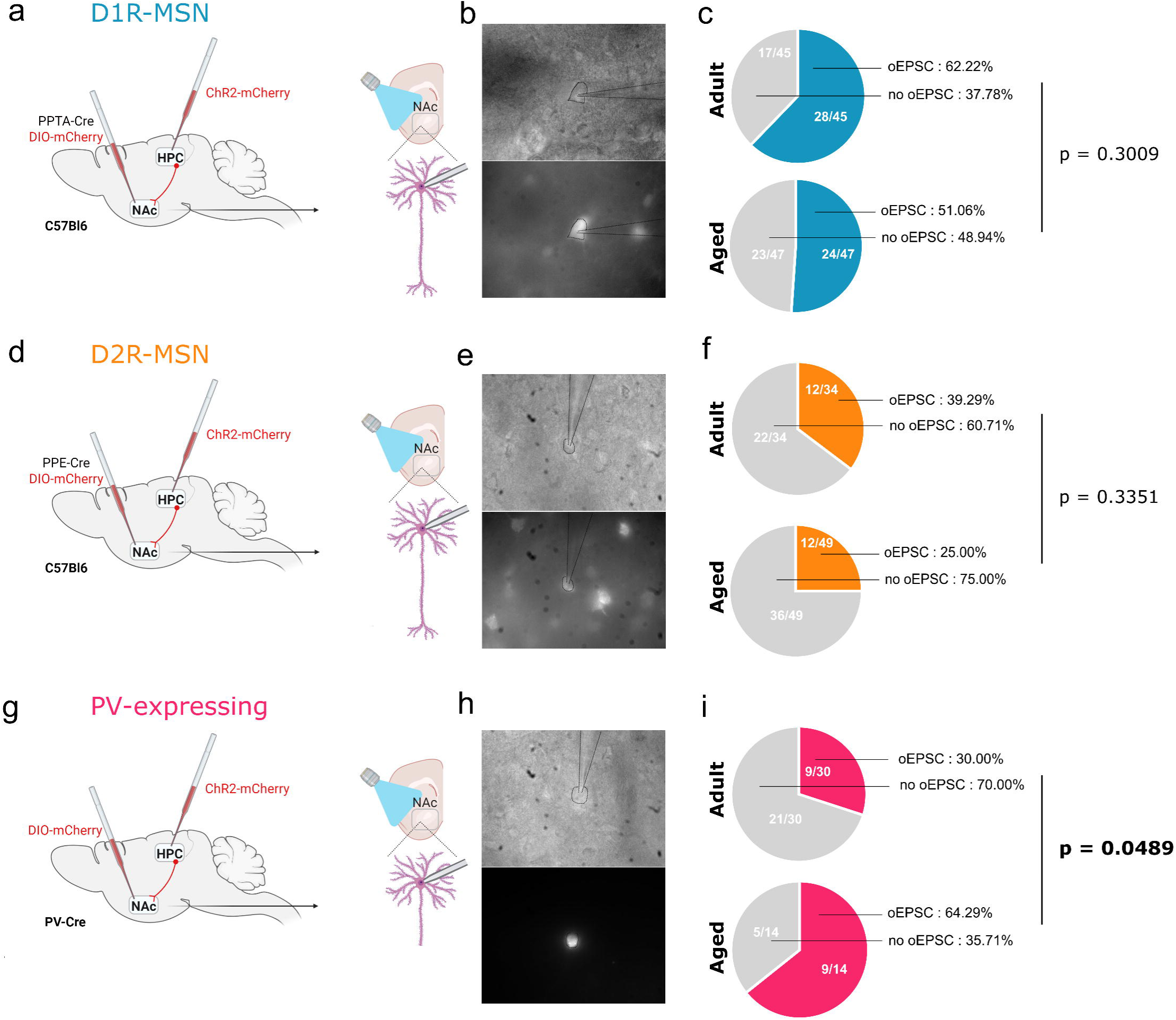
**Aging specifically alters dHPC**^◊**PV-expressing**^ **^neurons^ connectivity.** **(a)** Schematic representation of stereotaxic injections: viral tracing with PPTA-Cre combined with DIO-mCherry injected in the NAc to label specifically the D1R-MSN population and AAV-ChR2-mCherry in the dHPC. **(b)** Representative images of a D1R-MSN cell with a recording pipette (magnification 600x), without and with fluorescence. **(c)** The proportion of D1R-MSN that elicited electrophysiological response to the optical stimulation of the fibres coming from dHPC is maintained in aged animals. Results are expressed as percentage. (Fisher’s exact test, p = 0.3009, adult N = 13, n = 45, aged N = 17, n = 47). **(d)** Schematic representation of stereotaxic injections: viral tracing with PPE-Cre combined with DIO-mCherry injected in the NAc to label specifically the D2R-MSN population and AAV-ChR2-mCherry in the dHPC. **(e)** Representative images of a D2R-MSN cell with a recording pipette (magnification 600x), without and with fluorescence. **(f)** The proportions of D2R-MSN that elicited electrophysiological response to the optical stimulation of the fibres coming from dHPC are maintained in aging. Results are expressed as percentage. (Fisher’s exact test, p = 0.3351, adult N = 8, n = 34, aged N = 10, n = 49). **(g)** Schematic representation of stereotaxic injections: viral tracing with DIO-mCherry injected in the NAc of PV-Cre mice to label specifically the PV-expressing population and AAV-ChR2-mCherry in the dHPC. **(h)** Representative images of a PV-expressing cell with a recording pipette (magnification 600x), without and with fluorescence. **(i)** The proportion of PV-expressing neurons that elicited electrophysiological response to the optical stimulation of the fibres coming from dHPC is increased in aging. Results are expressed as percentage. (Fisher’s exact test, p = 0.0489, adult N = 7, n = 30, aged N = 4, n = 14). In all charts results are represented as number of cells with the indicated response/ total number of recorded cells.

### Aging oppositely modulates dHPC^→NAc^ synaptic strength onto D1R- vs D2R-MSNs

After determining dHPC^→NAc^connectivity, we next examined whether aging impacts dHPC^→NAc^synaptic strength (Fig. 4, Extended data Fig. 4). Evoked optogenetically excitatory postsynaptic currents (oEPSC, Fig. 4c), sensitive to the AMPA receptor antagonist, were observed (Fig. 4d), thus showing the glutamatergic nature of this synapse. The glutamatergic synaptic strength was significantly higher in aged animals (Fig. 4c). In order to investigate if dHPC^→NAc^could trigger inhibitory responses, we holded the same neurons at Vh=+10 mV. This allows to isolate GABAergic currents. We revealed a significant enhancement of inhibitory transmission in aged animals (Fig. 4e), sensitive to GABA-A receptor antagonist (Fig. 4f). As shown, only a subset of D1R-MSN shows both responses (Fig. 4g,h), a phenotype that is maintained with aging (Fig. 4h). In striking contrast, dHPC^→D2R-MSN^evoked responses showed an opposite profile to D1R-MSN, with the oEPSC amplitude showing a pronounced decrease with age (Fig. 4k,l), without changes in the evoked GABAergic transmission in the D2R-MSN subset showing both responses (Fig. 4m-o). Interestingly, we could observe D2R-MSN showing exclusively inhibitory responses, suggesting a highly regulated neuronal population within the NAc microcircuitry (Fig. 4p). Finally, we examined dHPC^→PV-expressing^neurons excitatory evoked transmission, in which we did not observe age-associated changes (Extended data Fig. 4). We rarely observed dHPC^→PV-expressing neurons^with both evoked excitatory and inhibitory responses. Aging oppositely alters dHPC^→NAc^glutamatergic synaptic strength: enhancement onto D1R-MSN and reduction onto D2R-MSN.

**Figure 4:**
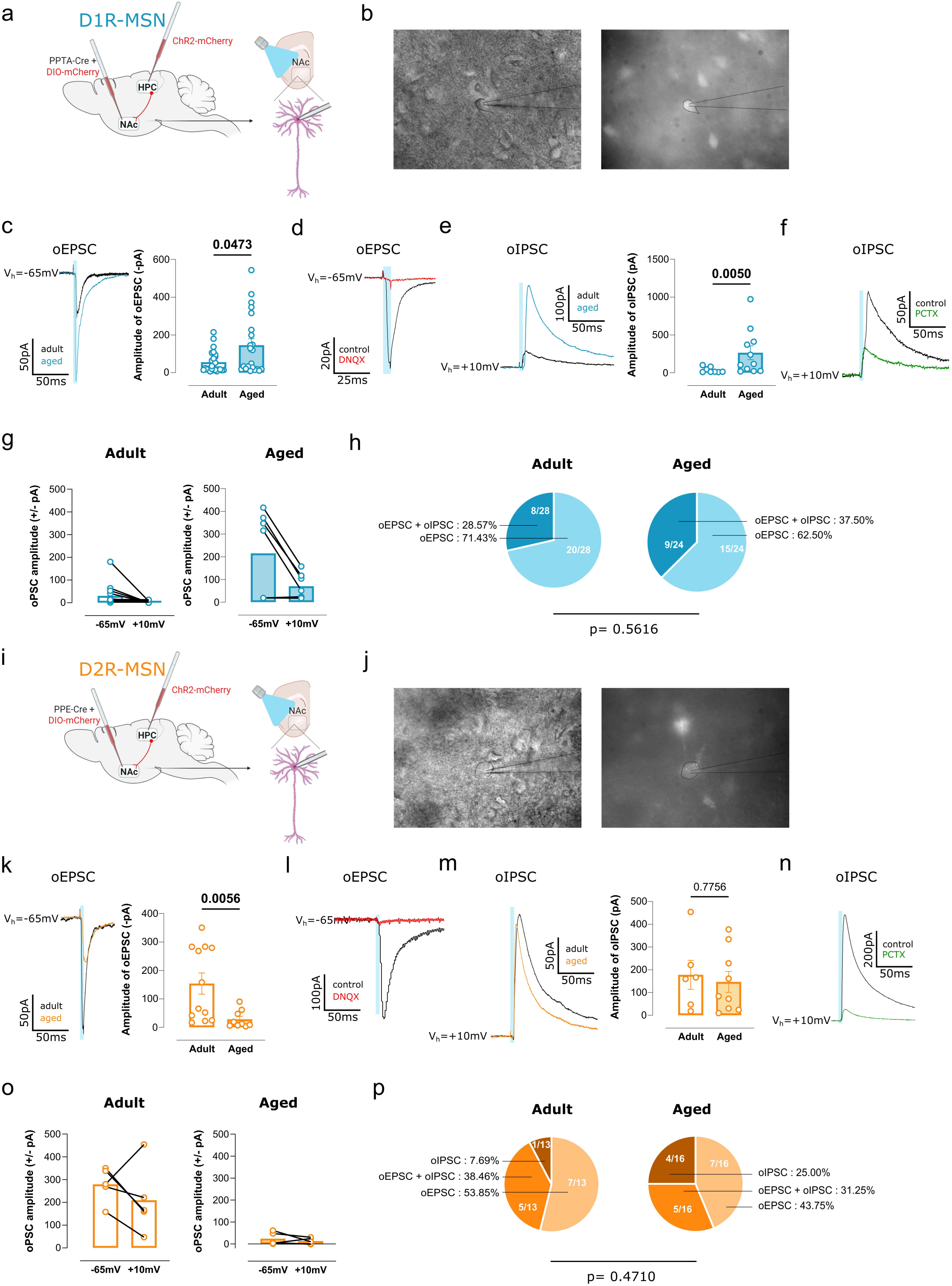
**Aging oppositely modulates dHPC**^◊**NAc**^ **synaptic strength onto D1R- vs D2R-MSNs.** **(a)** Schematic representation of stereotaxic injections: viral tracing with PPTA-Cre combined with DIO-mCherry injected in the NAc to label specifically the D1R-MSN population and AAV-ChR2-mCherry in the dHPC. **(b)** Representative images of a D1R-MSN cell with a recording pipette (magnification 600x), without and with fluorescence. **(c)** Representative traces of optogenetically-evoked excitatory post synaptic currents (oEPSC) after a 5ms blue light stimulation, represented as a blue bar. Amplitude of oEPSC is increased in aged D1R-MSN. Results are expressed as the mean ± SEM. (Normality test not passed. Mann-Whitney test, p = 0.0473, adult N = 11, n = 25, aged N = 12, n = 22). **(d)** Representative traces of optogenetic stimulation of the dHPC terminals eliciting oEPSC in D1R-MSN before (black) and after DNQX (red), a glutamatergic antagonist revealing the glutamatergic nature of the dHPC^◊D1R-MSN^projection. **(e)** Representative traces of optogenetically-evoked inhibitory post synaptic currents (oIPSC) after a 5ms blue light stimulation, represented as blue bar. Amplitude of oIPSC is increased in aged D1R-MSN. Results are expressed as the mean ± SEM. (Normality test not passed. Mann-Whitney test, p = 0.0050, adult N = 5, n = 8, aged N = 4, n = 11). **(f)** Representative traces of optogenetic stimulation of the dHPC terminals eliciting oIPSC in D1R-MSN before (black) and after PCTX (green), a GABA_A_ receptor antagonist revealing the GABAergic nature of the light evoked dHPC^◊D1R-MSN^response at V_h_ = +10mV. **(g)** Optogenetically-evoked post synaptic currents for the same cells at V_h_ = -65mV (excitatory) and V_h_ = -10mV (inhibitory). Results are expressed as the mean ± SEM. (For adult group: N = 5, n = 14. For aged group: N = 3, n = 7). **(h)** The proportion of D1R-MSN eliciting oEPSC or both oEPSC and oIPSC is maintained in aging. Results are expressed as percentages. (Fisher’s exact test, p = 0.5616, adult N = 11, n = 28, aged N = 12, n = 24). **(i)** Schematic representation of stereotaxic injections: viral tracing with PPE-Cre combined with DIO-mCherry injected in the NAc to label specifically the D2R-MSN population and AAV-ChR2-mCherry in the dHPC. **(j)** Representative images of a D2R-MSN cell with a recording pipette (magnification 600x), without and with fluorescence. **(k)** Representative traces of oEPSC after a 5ms blue light stimulation, represented as blue bar. Amplitude of oEPSC is decreased in aged D2R-MSN. Results are expressed as the mean ± SEM. (Normality test not passed. Mann-Whitney test, p = 0.0056, adult N = 6, n = 12, aged N = 7, n = 9). **(l)** Representative traces of optogenetic stimulation of the dHPC terminals eliciting oEPSC in D2R-MSN before (black) and after DNQX (red), a glutamatergic antagonist revealing the glutamatergic nature of the dHPC^◊D2R-MSN^projection. **(m)** Representative traces of oIPSC after a 5ms blue light stimulation, represented as blue bar. Amplitude of oIPSC is maintained for aged D2R-MSN. Results are expressed as the mean ± SEM. (Normality test not passed. Mann-Whitney test, p = 0.7756, adult N = 5, n = 6, aged N = 5, n = 9). **(n)** Representative traces of optogenetic stimulation of the dHPC terminals eliciting oIPSC in D2R-MSN before (black) and after PCTX (green), a GABA_A_ receptor antagonist revealing the GABAergic nature of the light evoked dHPC^◊D2R-MSN^response at V_h_ = +10mV. **(o)** Optogenetically-evoked post synaptic currents for the same cells at V_h_ = -65mV (excitatory) and V_h_ = -10mV (inhibitory). Results are expressed as the mean ± SEM. (For adult group: N = 4, n = 8. For aged group: N = 3, n = 5). **(p)** The proportion of D2R-MSN eliciting oEPSC or both oEPSC and oIPSC or oIPSC only is maintained in aging. Results are expressed as percentages. (Chi-square test, p = 0.4710, adult N = 8, n = 13 aged N = 10, n = 16). In all panels: significant p-values are represented in bold, N = number of animals, n = number of cells. In panels (h, p) results are represented as number of cells with the indicated response/ total number of recorded cells.

### Long-range dHPC-PV-expressing ^→NAc^ show a glutamatergic phenotype

As we observed optical evoked excitatory and inhibitory responses in both D1R-MSN and D2R-MSN, we hypothesized that the oIPSC were elicited by GABA released from the long-range dHPC-PV-expressing^→NAc^terminals. We thus compared the light elicited excitatory and inhibitory currents latency (Fig. 5) in order to discriminate direct from indirect synapses. Though neurons from both adult and ages animals exhibited similar kinetics (Fig. 5a-c, e-g, Extended data Fig. 5 a-c), D1R-MSN postsynaptic currents were time-locked to the optic stimulus, with oEPSC showing a fixed and short latency to the onset of optical stimulation (Fig. 5a,c). oIPSC displayed a delayed onset of approximately 5ms, indicative of an indirect polysynaptic pathway (Fig. 5b,c). Accordingly, blocking dHPC^→NAc^synaptic transmission with DNQX (6,7-dinotroquinoxaline-2,3-dione) abolished the optogenetically elicited GABAergic current (Fig. 5d), confirming its dependence on upstream glutamatergic activity and supporting the notion of an indirect connection. These effects were observed consistently across the different cell types herein investigated (Fig. 5e-h; Extended data Fig. 5). Our results suggest that the dHPC does not directly innervate the NAc with GABAergic inputs. Instead, a newly described long range dHPC PV-Glut co-expressing that project to the NAc is herein unveiled.

**Figure 5:**
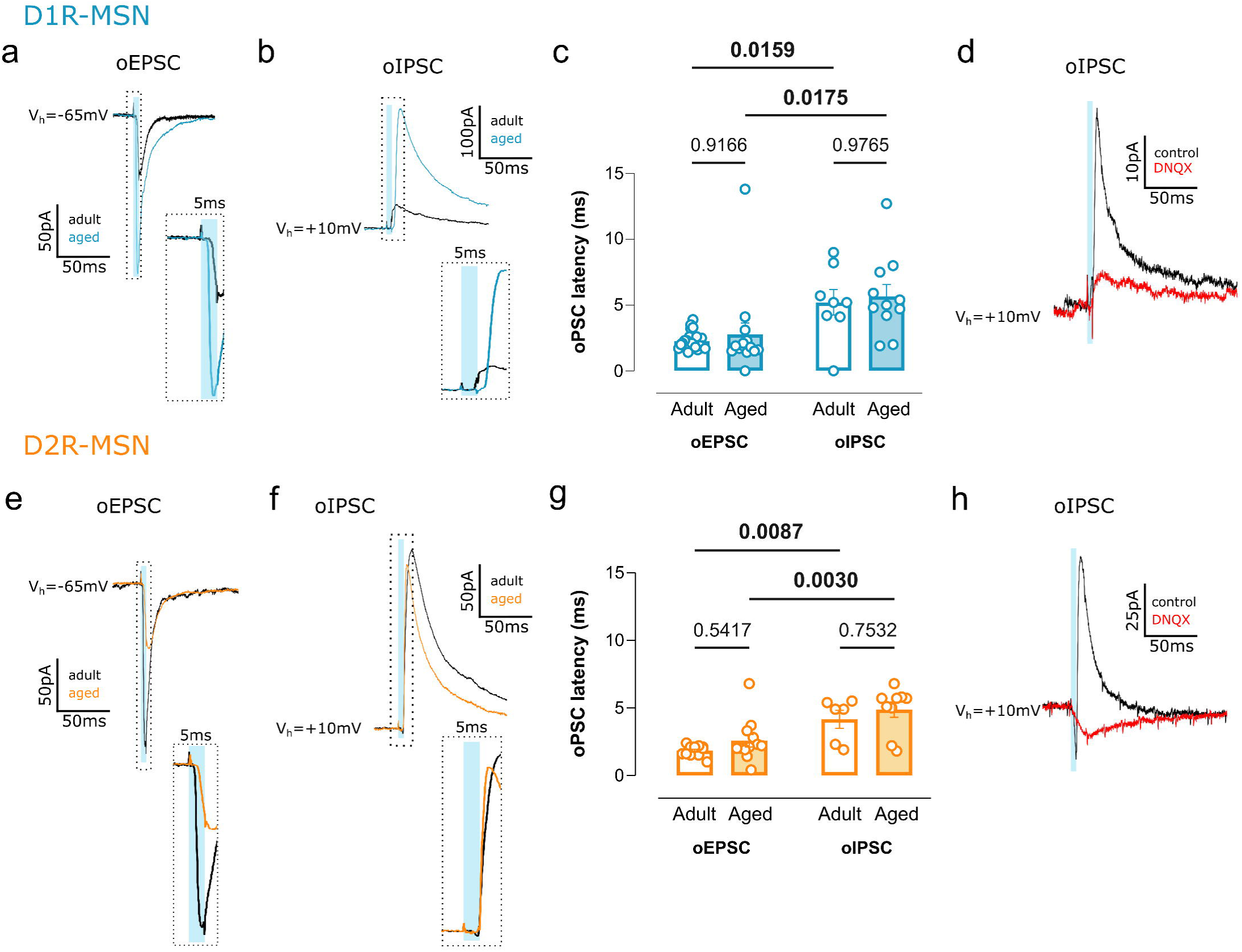
**Long-range dHPC PV-expressing**^◊**NAc**^ **show a glutamatergic phenotype.** **(a)** Representative traces of optogenetically-evoked excitatory post synaptic currents (oEPSC) after a 5ms blue light stimulation, represented as blue bar. Right: enlarged illustration of the dashed rectangle. **(b)** Representative traces of optogenetically-evoked inhibitory post synaptic currents (oIPSC) after a 5ms blue light stimulation, represented as blue bar. Right: enlarged illustration of the dashed rectangle. **(c)** Latency of oEPSC and oIPSC is preserved in aging for D1R-MSN, with oIPSC showing higher latency (around 5ms) compared to oEPSC (around 2ms). Results are expressed as the mean ± SEM. (Two-way ANOVA, V hold factor: p < 0.0001, F(1, 54) = 18.97, oEPSC adult N = 11, n = 25, aged N = 13, n = 14; oIPSC adult N = 5, n = 8, aged N = 4, n = 11). **(d)** Representative traces of optogenetic stimulation of the dHPC terminals eliciting oIPSC in D1R-MSN before (black) and after DNQX (red), a glutamatergic receptor antagonist revealing the dependence for the inhibitory transmission on upstream glutamatergic activity. **(e)** Representative traces of oEPSC after a 5ms blue light stimulation, represented as blue bar. Right: enlarged illustration of the dashed rectangle. **(f)** Representative traces of oIPSC after a 5ms blue light stimulation, represented as blue bar. Right: enlarged illustration of the dashed rectangle. **(g)** Latency of oEPSC and oIPSC is maintained in aging for D2R-MSN, with oIPSC showing higher latency (around 5ms) compared to oEPSC (around 2ms). Results are expressed as the mean ± SEM. (Two-way ANOVA, V hold factor: p < 0.0001 F(1, 35) = 25.68, oEPSC adult N = 6, n = 12, aged N = 7, n = 12; oIPSC adult N = 5, n = 6, aged N = 5, n = 9). **(h)** Representative traces of optogenetic stimulation of the dHPC terminals eliciting oIPSC in D2R-MSN before (black) and after DNQX (red), a glutamatergic receptor antagonist revealing the dependence for the inhibitory transmission on upstream glutamatergic activity. In all panels: significant p-values are represented in bold, N = number of animals, n = number of cells.

### The age-related changes in the dHPC^→NAc^ display a different pattern in females

Cognitive decline has been shown to exhibit sex specificities^7^. Also, some studies report that the hippocampal volume decline is more evident in men from 50 years old onwards^24,25^. We therefore investigated if the age-associated changes unveiled in males above, could also be observed in females. Spontaneous activity analysis (Extended Fig. 6,7,8) shows that in aging D2R-MSN present altered E/I ratio towards inhibition (Extended Fig. 7n), which results from enhanced amplitude and frequency of spontaneous inhibitory inputs (Extended Fig. 7i-l), thus suggesting that in females the NAc microcircuitry is differently affected by aging when compared to males. Interestingly, unlike males, a similar proportion within the different NAc neuron populations is connected to the dHPC, i.e., approximately 2/3, and this remains unaltered with aging. Evoked light activation of the dHPC^→NAc^pathway resulted in a tendency to enhanced synaptic strength in both MSN populations whereas opposite trend was observed in PV-expressing neurons (Fig. 6d,j,i). As in males, we assessed light evoked excitatory and inhibitory responses in the same neuron. A higher proportion of D1R-MSN displayed both responses, whereas light-evoked inhibitory responses were scarcer in D2R-MSN and PV-expressing neurons (Fig. 6 f,k,p). As in males, these responses showed increased latency (Extended data Fig. 9). Thus, females show distinct patterns of age-related alterations in the dHPC^→NAc^pathway in comparison with males, thus suggesting structural and functional sex specificities.

**Figure 6:**
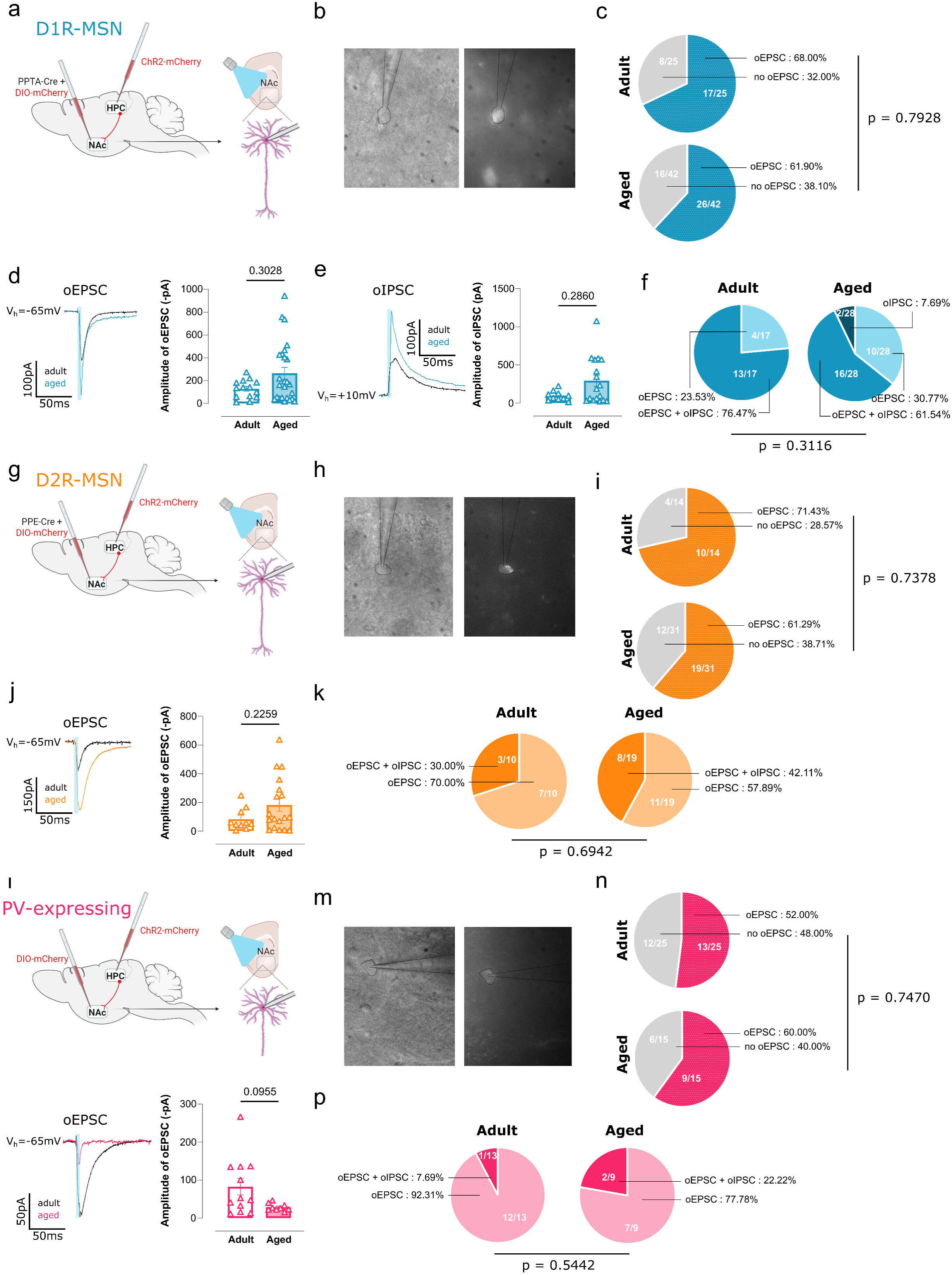
**The age-related changes in the dHPC**^◊**NAc**^ **display a different pattern in females.** **(a)** Schematic representation of stereotaxic injections: viral tracing with PPTA-Cre combined with DIO-mCherry injected in the NAc to label specifically the D1R-MSN population and AAV-ChR2-mCherry in the dHPC in females. **(b)** Representative images of a D1R-MSN cell with a recording pipette, without and with fluorescence (magnification 600x). **(c)** Optical stimulation of dHPC^◊NAc^fibres elicited D1R-MSN responses with preserved connectivity. Results are expressed as percentage. (Fisher’s exact test, p = 0.7928, adult N = 9, n = 25, aged N = 12, n = 42). **(d)** Representative traces of optogenetically-evoked excitatory post synaptic currents (oEPSC) after a 5ms blue light stimulation, represented as blue bar. Amplitude of oEPSC is maintained in aged D1R-MSN. Results are expressed as the mean ± SEM. (Normality test not passed. Mann-Whitney test, p = 0.3028, adult N = 8, n = 14, aged N = 11, n = 25). **(e)** Representative traces of optogenetically-evoked inhibitory post synaptic currents (oIPSC) after a 5ms blue light stimulation, represented as blue bar. Amplitude of oIPSC is maintained in aged D1R-MSN. Results are expressed as the mean ± SEM. (Normality test not passed. Mann-Whitney test, p = 0.2860, adult N = 8, n = 10, aged N = 11, n = 17). **(f)** The proportion of D1R-MSN eliciting oEPSC, both oEPSC and oIPSC or oIPSC only is maintained in aging. Results are expressed as percentages. (Fisher’s exact test, p = 0,3116, adult N = 8, n = 17, aged N = 11, n = 28). **(g)** Schematic representation of stereotaxic injections: viral tracing with PPE-Cre combined with DIO-mCherry injected in the NAc to label specifically the D2R-MSN population and AAV-ChR2-mCherry in the dHPC in females. **(h)** Representative images of a D2R-MSN cell with a recording pipette, without and with fluorescence (magnification 600x). **(i)** Optical stimulation of dHPC^◊NAc^fibres elicited D2R-MSN responses with preserved connectivity. Results are expressed as percentage. (Fisher’s exact test, p = 0.7378, adult N = 3, n = 14, aged N = 7, n = 31). **(j)** Representative traces of oEPSC after a 5ms blue light stimulation, represented as blue bar. Amplitude of oEPSC is preserved in aged D2R-MSN in females. Results are expressed as the mean ± SEM. (Normality test not passed. Mann-Whitney test, p = 0.2259, adult N = 3, n = 10, aged N = 7, n = 18). **(k)** The proportion of D2R-MSN eliciting oEPSC or both oEPSC and oIPSC is maintained in aging. Results are expressed as percentages. (Fisher’s exact test, p = 0.6942, adult N = 8, n = 10, aged N = 11, n = 19). **(l)** Schematic representation of stereotaxic injections: viral tracing with DIO-mCherry injected in the NAc to label specifically the PV-expressing neuronal population and AAV-ChR2-mCherry in the dHPC in females. **(m)** Representative images of a PV-expressing cell with a recording pipette, without and with fluorescence. **(n)** Optical stimulation of dHPC^◊NAc^fibres elicited PV-expressing neuron responses with preserved connectivity. Results are expressed as percentage. (Fisher’s exact test, p = 0.7470, adult N = 5, n = 25, aged N = 5, n = 15). **(o)** Representative traces of oEPSC after a 5ms blue light stimulation, represented as blue bar. Amplitude of oEPSC is maintained in aged PV-expressing neurons in females. Results are expressed as the mean ± SEM. (Normality test not passed. Mann-Whitney test, p = 0.0955, adult N = 5, n = 12, aged N = 5, n = 9). **(p)** The proportion of PV-expressing cells eliciting oEPSC or both oEPSC and oIPSC is maintained in aging. Results are expressed as percentage. (Fisher’s exact test, p = 0,5442, adult N = 7, n = 13, aged N = 4, n = 9). In all panels: significant p-values are represented in bold, N = number of animals, n = number of cells.

### Reducing activity in the dHPC^→NAc^ circuit improves the reward-driven spatial memory specifically in aged mice

Aging is characterized by cognitive heterogeneity, potentially linked to the cellular alterations in the dHPC^→NAc^pathway. To test this, we carried out behavioral analysis, including a reward-driven spatial memory task, known to implicate the dHPC^→NAc^pathway^40^ (Fig. 7a,b). In average, all groups were able to discriminate the compartment associated to the reward. Yet, aged mice (both in males and females) showed increased variance with a trajectory towards weaker reward-driven spatial memory (Fig. 7c,d). These results strengths this age as a critical window in which divergent trajectories towards cognitive decline start. Anxiety levels and hedonic responses were preserved in aged mice (Fig. 7e,f).

**Figure 7:**
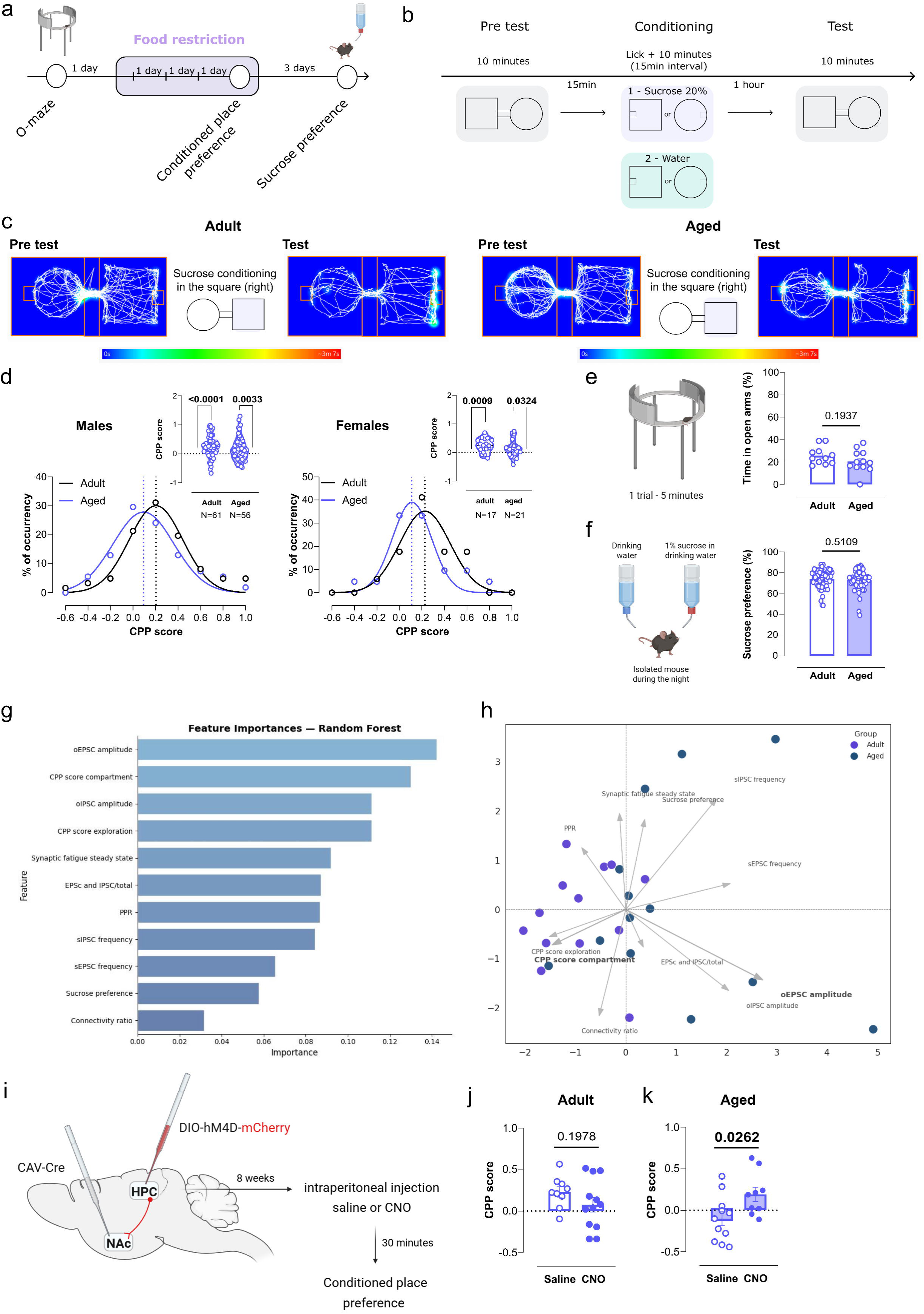
**Reducing activity in the dHPC**^◊**NAc**^ **circuit restores the reward-driven spatial memory in aged mice** **(a)** Behavioral assessment experimental design. **(b)** Diagram of conditioned place preference protocol (CPP). **(c)** Representative heat maps and locomotion tracking of the pre-test and test phases. The scale represents the average time spent in a specific location. The tracking follows the head of the animal. A more pronounced preference for the compartment associated with the reward is observed during the test for the adult mouse. **(d)** Distribution of the CPP score for males (left) and females (right) showing a shift towards weaker reward-driven spatial memory for aged animals (Males: adult N = 61, aged N = 56; Females: adult N = 17, aged N = 21). On top of each distribution graph, the CPP score is shown and results are expressed as the mean ± SEM. (Males: One sample t-test, p < 0.0001 for adult N = 61, p = 0.0033 for aged N = 56; Females: One Sample t-test, p = 0.0009 for adult N = 17, p = 0.0324 for aged N = 21). **(e)** Anxiety levels are preserved with aging, as both groups presented similar results for the zero-maze test. Results are expressed as the mean ± SEM. (Passed normality test. Unpaired t-test, p = 0.1937, adult N = 11, aged N = 12). **(f)** The hedonic response was also maintained with aging as the sucrose preference test showed similar results in adult and aged mice. Results are expressed as the mean ± SEM. (Normality test not passed. Mann-Whitney test, p = 0.5109, adult N = 55, aged N = 63). **(g)** Random forrest analysis identifying the top predictive behavioral and electrophysiological variables distinguishing adult and aged mice (top features explained > 60% of classification accuracy by cumulative importance. (Feature importance values shown as % of mean decrease in Gini impurity across tress.) **(h)** Principal component analysis (PCA) showing age-dependant separation along PC1 and PC2, with the two components capturing around 45-55% of the total variance in behavioral and synaptic metrics. **(i)** Schematic representation of stereotaxic injections and timeline: intersectional viral tracing with a retrograde CAV2-Cre injected in the NAc and AAV-hM4D-mCherry in the dHPC to specially hyperpolarize *in vivo* the dHPC neurons projecting to the NAc, thus reducing their activity. Clozapine-N-oxide (CNO) was injected 30 minutes before the CPP. **(j)** Reducing activity of the dHPC^◊NAc^projection by CNO treatment improved the reward-driven spatial memory, reported as CPP score, specifically in aged animals. Results are expressed as the mean ± SEM. (Adult: Passed normality test. Unpaired t-test, p = 0.1978, adult control N = 9 / CNO N = 12. Aged: Passed normality test. Unpaired t-test, p = 0.0262, aged control N = 11 / CNO N = 9). In all panels: significant p-values are represented in bold, N = number of animals.

To identify which age-related changes best account for the differences observed in aged mice, we performed a random forest analysis using the variables collected for each animal. We first applied principal component analysis (PCA) to the combined dataset comprising both D1R-and D2R-MSN. Because this initial analysis revealed a clearer separation by age within the D1R-MSN population, subsequent PCA was restricted to the D1R-MSN subset. On this basis, we generated a D1R-MSN mouse pipeline matrix. Adult and aged mice showed a significant variance, with two main contributing factors being: oEPSC amplitude and conditioned place preference (CPP) score (Fig. 7g). Principal component analysis (Fig. 7h) further illustrated the variability among aged animals, indicating that higher synaptic strength correlated with poorer reward-driven spatial memory. To test whether a negative behavioral trajectory in aged mice could be causally linked to cellular changes, we used *in vivo* chemogenetics. We injected a retrograde CAV2-Cre virus into the NAc and a Cre-dependent hM4D receptor (AAV-DIO-hM4-mCherry) in the dHPC to specifically induce the hypoactivation of dHPC^→NAc^projecting neurons (Fig. 7i). Aged mice treated with clozapine-N-oxide (CNO) showed improved CPP score, thus corroborating our hypothesis that dHPC^→NAc^hyperactivity is deleterious for the reward-driven spatial memory. CNO treatment in adult mice perturbed appetitive memory, as shown by others^40^ (Fig. 7j). These findings show that dHPC^→NAc^pathway finely tunes reward-driven spatial memory following a bell-shape profile.

### Human fMRI reveals increased hippocampal activity and strengthened posterior HPC-NAc correlation during navigation encoding in healthy older participants

To translate our rodent findings to humans, we examined human fMRI data acquired during a passive navigation task in young and healthy older participants (Fig. 8a). The present neuroimaging study is a re-analysis of functional data (Fig. 8b) from twenty-five younger (∼28 years old) and twenty-six older healthy adults (∼73 years old) from the French longitudinal cohort study Silver Sight^45^ performing a two-choices navigation task^46^, fully described in the methods. The linear model revealed main effects of group (F(1,195) = 18.657, p < 0.001, partial η2 = 0.087), ROI (F(2,195) = 4.620, p = 0.011, partial η2 = 0.045), and contrast (F(1,195) = 4.040, p = 0.046, partial η2 = 0.020) factors on parameter estimates. Post hoc tests showed that during the Encoding condition older participants exhibited significantly greater brain activity than young group in the anterior HPC (aHPC) (Fig. 8c) (t(34) = 3.96, p = 0.00036, 95% CI [0.80, 2.49], SE = 0.42) and posterior HPC (pHPC) (t(35) = 2.29, p = 0.028, 95% CI [0.17, 2.73], SE = 0.63), but there was no significant age group effect for the NAc (t(32) = 1.34, p = 0.190, 95% CI [-0.56, 2.73], SE = 0.81]. Finally, we conducted Pearson correlations between HPC and NAc parameter estimates to examine HPC-NAc dynamic. Significant positive correlation was found in the older group during the Encoding phase between pHPC and NAc (Fig. 8d) (N = 14) (r= 0.64, p= 0.014). We did not observe significant pHPC-NAc correlations among young participants (N = 20) (r = -0.24, p = 0.30). Overall, these findings indicate a shift toward increased hippocampal activity in older participants during the navigation task, with a significant positive pHPC-NAc correlation during the encoding phase, uniquely present in this group, mirroring our observations in rodents.

**Figure 8:**
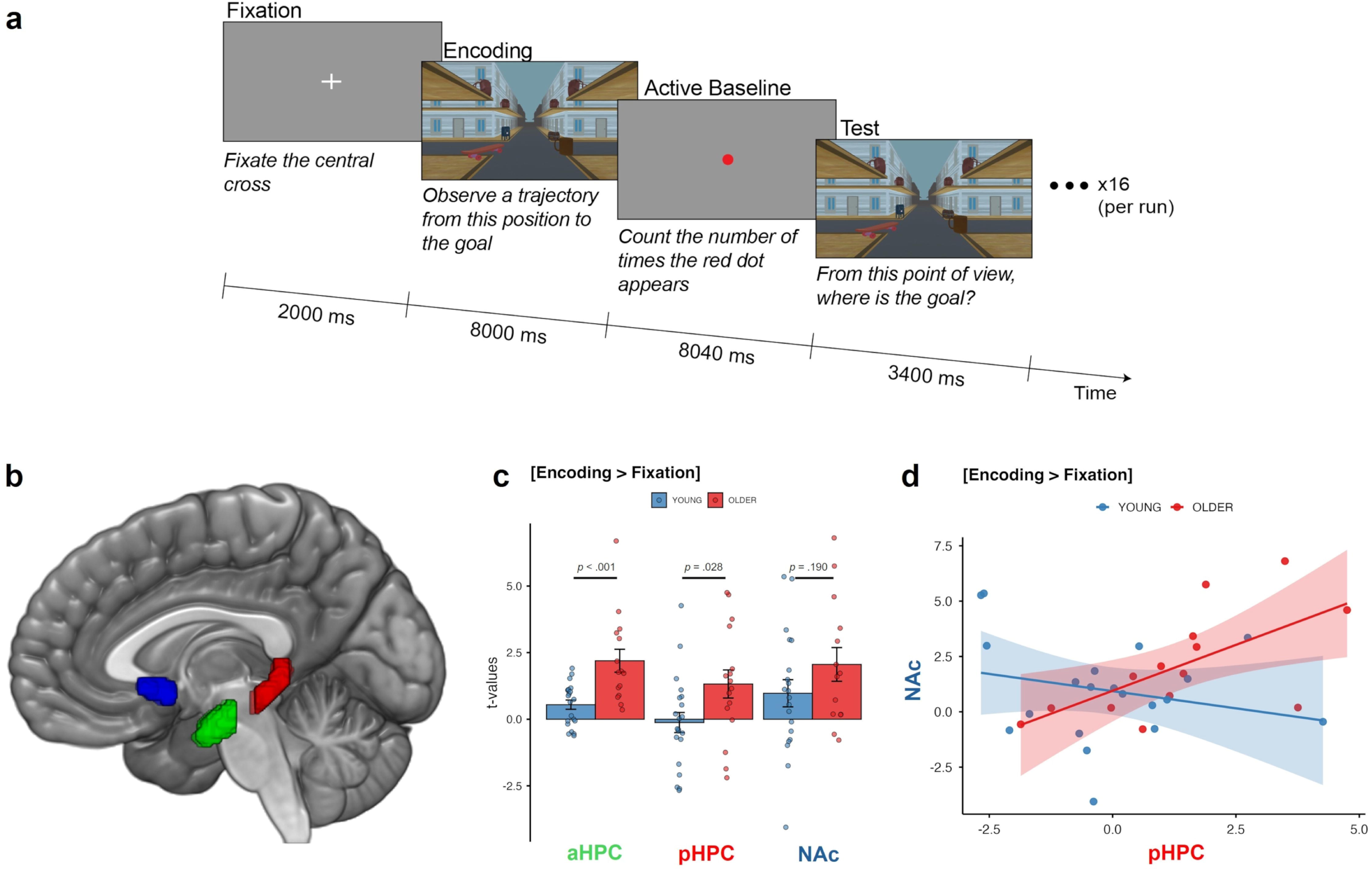
fMRI reveals increased hippocampal activity and strengthened pHPC–NAc correlation during navigation encoding in older participants (a) Schematic representation of a single trial within a run. There are 16 trials in a run. Participants first fixate a central cross. They observe a trajectory through an environment with a unique object configuration and they learn the position of the goal. Then, they count the number of times a red dot flashes during the active baseline task. Finally, they are put back into the previously learned environment in front of the intersection and they need to decide if the goal is located to their right or to their left. **(b)** Regions of Interests (ROIs) with nucleus accumbens (blue, Nac), anterior hippocampus (green, aHPC), and posterior hippocampus (red, pHPC) are shown on a 3D-rendered brain. **(c)** Brain activity the Encoding phase compared to the Fixation condition [Encoding > Fixation] for young (blue) and older adults (red) in each ROIs. **(d)** Pearson correlations patterns between brain activity in the nucleus accumbens (NAc) and the posterior hippocampus (pHPC) during encoding phase compared to the Fixation condition [Encoding > Fixation] for young (blue) and older adults (red).

## Discussion

Our findings reveal that aging is associated with a shift in the E/I balance toward increased excitation in dHPC CA1 pyramidal neurons, generating a hyperactive state in this major hippocampal output region. Within the dHPC^→NAc^pathway, we identify two neuronal populations - regular-spiking and fast-spiking - with distinct electrophysiological profiles, including a previously undescribed long-range PV-Glut co-expressing fast-spiking population. We further show that the dHPC^→NAc^monosynaptic pathway differentially modulates synaptic strength onto D1R- and D2R-MSN, with enhanced inputs onto D1R-MSN negatively associated with reward-driven spatial memory performance. Selective chemogenetic reduction of hyperactivity in dHPC^→NAc^projecting neurons improves reward-driven spatial memory specifically in aged mice. Of interest, older healthy human participants performing a goal-directed virtual navigation task exhibit heightened posterior hippocampal activity that correlates with increased NAc activity during the encoding phase. Together, these results show that HPC^→NAc^pathway reinforcement marks a shift towards cognitive decline. Thus, targeted modulation of dHPC^→NAc^connectivity may offer new strategies to maintain cognitive function and correct negative cognitive trajectories.

Hippocampal volume decreases with aging, which has been linked to reduced white matter^47^ and synaptic loss^48,49^. One might hypothesize reduced hippocampal neuronal activity and connectivity as a result. Yet, the present work demonstrates hippocampal hyperactivity in CA1 neurons at early stages of aging in mice, arising from an increased E/I balance. In humans, older individuals similarly show increased hippocampal fMRI signals during a navigation task, indicating convergent evidence across species. Two conceptual frameworks co-exist to explain this hyperactivity: one, it acts as a compensatory response to emerging synaptic and connectivity dysfunction^50^, second it is maladaptive, promoting network instability^51^, but causality proof remains sparse. Our data support hippocampal hyperactivity as a detrimental mechanism, reflected in maladaptive HPC-NAc coupling negatively associated with reward-driven spatial memory, which can be alleviated by fine-tuned reduction of dHPC^→NAc^ excitability. Studies in the literature, in both humans and rodents, align with our data, showing hyperactivity at early stages of aging^52–55^. Here, we demonstrate that hippocampal hyperactivity functionally impacts an aging-sensitive target region, identifying HPC^→NAc^reinforcement as a biomarker of cognitive vulnerability that is amenable to intervention at an early stage.

We herein describe that dHPC projecting to the NAc show two distinguished electrophysiological profiles: neurons exhibiting the canonical firing properties of CA1 pyramidal glutamatergic neurons, consistent with established literature^56^, and a population with a fast-spiking profile reminiscent of PV-expressing interneurons in the dHPC^57^. We gathered strong evidence by patch-clamp electrophysiology and immunohistochemistry that PV-expressing neurons in the dHPC, both in the pyramidal layer and stratum oriens are long-range projecting neurons. We now show evidence for a dHPC PV-Glut co-expressing neurons projecting to the NAc, as the light evoked inhibitory responses recorded in the NAc presented a delay and were sensitive to glutamatergic receptor antagonist. Occasional reports have shown that PV can be co-expressed with the glutamate transporters Vglut1 or Vglut2 in neurons from different regions of the adult rodent’s brain: lateral hypothalamus^58,59^, anterior hypothalamus^60^, lateral habenula^61^ and distal subicculum of the HPC^62^. These findings have gained support from single cell transcriptome analysis of brain neurons^63^. Of note, the excitability profile of long-range PV-expressing-glutamatergic neurons reported in Brenton, 2023^60^ is similar to the one we herein report.

In males, the synaptic strength of dHPC inputs onto D1R-MSN and D2R-MSN in the NAc undergoes distinct age-dependent changes, leading to a shift towards enhanced D1R-MSN reinforcement accompanied by a weakening of D2R-MSN activity. We also observed increased connectivity onto PV-expressing neurons, which may reflect a homeostatic plasticity mechanism compensating for the heightened dHPC^→D1R-MSN^synaptic strength. Extensive work in adult rodents shows that D1R- and D2R-MSN have complementary roles in reward, motivation, and cognition. Their activity reliably encodes positive and negative valence, and both populations form similar functional clusters during Pavlovian conditioning, jointly supporting associative learning^64^. D1R-MSN respond to salient stimuli, whereas D2R-MSN signal prediction errors^65^. Moreover, D2R-MSN hypoactivity increases inappropriate behaviours, while their activation enhances reversal learning^66^. In aging, the expression of D2 dopamine receptors declines^67,68^, as do dopamine levels in the NAc^69^ in both rodents and humans. This could account for our observation that dHPC^→D2R-MSN^synapses exhibit weakened responses in the aged NAc. dHPC^→NAc^synaptic strength shifts from a D2R-MSN > D1R-MSN balance in adults to a D1R-MSN > D2R-MSN dominance in aged mice, perturbing the animals’ ability to associate a context with a pre-exposed natural reward. This occurs without altering the E/I balance within the NAc microcircuit of the cell populations examined, nor NAc-related behaviors such as hedonia. Together, these findings strongly suggest that the dHPC^→NAc^pathway exhibits early vulnerability prior to the emergence of detrimental changes within the NAc itself.

Our study reveals pronounced sex differences in the vulnerability of the NAc microcircuitry and HPC^→NAc^pathway to aging. These differences raise the question of whether females engage similar neuronal circuits and cellular processes to support reward driven associative behaviours. In this line, studies demonstrate that males and females recruit similar brain regions during spatial tasks but exhibit distinct activation intensities, explaining divergent neural signatures despite overlapping functionality^70,71^. Hippocampal aging trajectories further highlight sexual dimorphism. A longitudinal study reports that over two years, women increase bilateral hippocampal connectivity, whereas men enhance connectivity between hippocampus and anterior cingulate cortex, indicating sex-specific circuit remodelling with age^72^. Recent work directly probing the HPC^→NAc^pathway reveals that, although males and females exhibit comparable synaptic plasticity, they rely on distinct molecular mechanisms: males depend on glutamatergic receptor activation, whereas females recruit voltage-gated calcium channels and estrogen receptor-α. These convergent yet mechanistically divergent pathways highlight how similar circuit-level outcomes can emerge from sexually dimorphic molecular processes^73^. In sum, our findings, together with prior literature, underscore the critical need to perform studies specifically in females. Multiple lines of evidence - including our own - demonstrate pronounced sex dimorphism, particularly at the cellular level.

By defining a targetable hippocampus–accumbens mechanism engaged in an early intervention-sensitive window, our findings open therapeutical avenues to recalibrate circuit dynamics and slow maladaptive cognitive trajectories with aging.

## Material and methods

### Animals

All procedures were in accordance with the recommendations of the European Commission (2010/63/EU) for care and use of laboratory animals and approved by the French National Ethical Committee (#40372-2022113017036060). We used male and female wild-type C57Bl6 mice and PV-Cre mice. Transgenic mice were heterozygous and backcrossed on a C57BL/6J background. They were housed 4-5 per cage under standard conditions, at 22°C, 55% to 65% humidity, with a 12-hour light/dark cycle (7am/7pm). Food and water were *ad libitum*, except for the food restriction 3 days before the conditioned place preference test. Enriched housing consisted of a chewing block of wood, a plastic igloo and cotton to facilitate nesting. Mice were used at different ages: adult (8-12 weeks) and aged (18-20 months).

### fMRI

#### Participants

The present neuroimaging study is a re-analysis of functional data from twenty-five younger and twenty-six older healthy adults from the French longitudinal cohort study *SilverSight*^45^ performing a two-choices navigation task^46^. It should be noted that this initial study focused exclusively on the brain activity of visual regions. Six older adult participants were excluded due to poor task understanding, and one young adult was excluded due to somnolence during the fMRI scanning. After the data preprocessing procedure, three older subjects were excluded due to excessive motion. Finally, one older and three young participants were removed from the analysis because of the missing runs or scans issue. The final sample consisted of twenty-one young adults (M = 28.05 years, SD = 3.75 years; 12 female) and sixteen older adults (M = 73.63 years, SD = 5.61 years; 7 female). All participants reported no history of neurological or psychiatric disorders. They provided written informed consent, and the study protocol was approved by the Ethics Committee “CPP ^Ile-de-France V” (ID RCB: 2015-A01094-45, CPP No: 16122).

#### Task

The virtual navigation task was developed using Unity3D v2019.2 (Unity Technologies, Inc. San Francisco, CA, USA) and displayed on an MRI-compatible liquid crystal display monitor (NordicNeuro-Lab, Bergen, Norway) positioned at the head of the scanner bore. Virtual environment depicted a four-way intersection from which started four streets lined with identical buildings and sidewalks, making them strictly indistinguishable from each other. The first floor of each building comprised a balcony. At the level of the intersection, the sidewalks and balconies formed 8 right angles in total onto which landmarks could be positioned. Each trial was characterized by a unique version of the intersection, and it consisted of 4 distinct phases. First, participants fixated a central white cross on a grey background for 2000 ms. Second, they performed the encoding phase of the navigation task. The latter projected participants into a first-person perspective and required them to observe a trajectory within the virtual environment. They started at the level of the intersection where they stayed for 3000 ms to allow sufficient time for item encoding. Then, the trajectory resumed and made a left or right turn into the street. The trajectory stopped upon reaching the goal (i.e., a flower bouquet), 10 virtual meters into the street. Finally, they performed the test phase of the navigation task. They were re-projected into a first-person perspective of the environment, facing the intersection either from the same learned direction or from the opposite direction. They had 3400 ms to decide whether the goal was situated in the street to their right or to their left by pressing a key using MRI-compatible ergonomic two-grip response device (NordicNeuroLab, Bergen, Norway). We did not provide participants with any feedback. Participants completed a total of 8 runs, each comprised of 16 trials. Each run included both original and flipped versions of the intersections, and half of the trials in each run involved testing participants from the viewpoint opposite to that used during the encoding phase. Run order was counterbalanced across participants using a balanced Latin square design.

#### MRI acquisition

Neuroimaging data were acquired using a 3-Tesla MAGNETOM Prisma MRI scanner (Siemens Medical Solution, Erlangen, Germany) equipped with a 64-channel head coil at the Neuroimaging and Radiology Unit of the Quinze-Vingts National Hospital in Paris. A total of 138 whole-brain volumes were obtained using T2*-weighted echo-planar imaging (EPI) sequences with the following parameters: voxel size = 3 × 3 × 2 mm3; repetition time (TR) = 2680 ms; echo time (TE) = 30 ms; flip angle = 90°; slice order = interleaved; number of slices = 48; matrix size = 74×74; field of view (FoV) = 220×220 mm. A structural 3D brain image was acquired using a T1-weighted high-resolution Magnetization Prepared Rapid Gradient Echo (MPRAGE) sequence with parameters: voxel size = 1 × 1 × 1.1 mm3; TR = 2300 ms; TE = 2.98 ms; flip angle = 9°; number of slices = 176; matrix size =256 × 248.

#### MRI preprocessing

The acquired fMRI data were analyzed using SPM12 (Wellcome Department of Imaging Neuroscience, London, UK) and custom codes in MATLAB R2024b. To ensure state equilibrium, the first and last five functional volumes were excluded from each run. The preprocessing pipeline followed standard steps. Slice-timing correction was applied to account for the interleaved order of slices. Realignment was used to correct for head motion by aligning all functional images to a mean image in each run. Six head movement parameters estimated in this step were further analyzed for motion artifacts by computing framewise displacement (FD) - the sum of the absolute temporal derivatives of the motion parameters. Volumes with FD > 0.9 mm were flagged as potential motion outlier^74,75^, and three older participants with more than 30% of flagged volumes in at least one of the runs were excluded from further analysis. The next steps involved co-registration of functional images to the participant’s structural image, normalization in order to align data into standard Montreal Neurological Institute (MNI) space, and smoothing to enhance signal-to-noise ratio with an 8 mm full-width at half maximum (FWHM) Gaussian kernel.

#### Regions of interest (ROIs)

ROIs included the anterior and posterior parts of hippocampus (aHPC and pHPC) as well as the Nucleus Accumbens (NAc) (Figure 8b). ROI masks for both hemispheres were extracted using the FSLeyes version 1.14.2 (McCarthy, 2025; University of Oxford, Oxford, UK) with the Harvard-Oxford subcortical structural atlases aligned to the MNI152 standard space^76^. The HPC in each hemisphere was segmented into anterior and posterior segments along its long axis based on an anatomical landmark of the uncal apex at y = −21 mm in MNI space^77^, corresponding to the voxel index y = 52. To prevent an overlap between the anterior and posterior HPC, a slice of 2 voxels was removed from each of the adjacent boundaries, leaving a gap zone of 4 voxels between the regions.

### Statistical Analyses

We fit the preprocessed data to subject-level GLMs with 18 regressors of interest for each run: (1) onsets and durations of the encoding phases from the 16 trials, (2) onsets and durations of the active baseline tasks from the 16 trials and (3) onsets and durations of the fixation phases from the 16 trials. We included the 3 rotation and 3 translation movement parameters as confound regressors. We also added a volume-wise regressor that modelled out volumes that were flagged during motion scrubbing. We convolved the GLMs with the SPM canonical hemodynamic response function (HRF) and we high-pass filtered (1/128 Hz cut-off) the time series from each voxel to remove low-frequency noise and signal drift. In order to elucidate the patterns of activity within the pHPC, aHPC and NAc in relation to the experimental conditions, we extracted t-activation maps from the individual ROI masks for the contrasts of interest [Encoding > Fixation] and [Test > Fixation] across age groups. First, we implemented linear mixed models to examine the effects of age group, ROI and condition on the t-maps associated with the 2 main contrasts including participants as a random intercept in all models. Post-hoc analysis involved two-way interactions, Main Effects, and Multiple Pairwise Comparisons tests. Outliers were removed from the fMRI parameters dataset to ensure the normality assumption of the residuals. A non-parametric boxplot rule was used to detect outliers: any data point lying outside the interval [Q1 −1.5 ・ IQR,Q3 +1.5 ・ IQR] was considered extreme, where Q1 and Q3 represent the first and third quartiles, respectively, and IQR = Q3 − Q1 (R Core Team, 2025, https://ropensci.org/blog/2021/11/16/how-to-cite-r-and-r-packages/).

Second, Behavioral data, including the percentage of correct responses and average reaction times (RT) across all 128 trials during the test phases of the virtual spatial task, were analyzed using Python 3.13.5 in Jupyter Notebook (version 7.4.4). Since the behavioral measures did not follow the normality assumption, non-parametric Spearman rank correlation tests were conducted in R to assess whether HPC and NAc activation during the contrasts of interest [Encoding > Fixation] and [Test > Fixation] in both young and older groups were associated with the behavioral performance. To study the anterior/posterior HPC-NAc functional activity, Pearson correlation coefficients were computed between the extracted contrast estimates for each group from pHPC, aHPC and NAc. Significance level threshold was set at p < 0.05/3 = 0.017 using Bonferroni correction to adjust for multiple comparisons across the three ROIs.

### Drug administration

Clozapine N-Oxyde was purchased from Enzo Life (France). The final concentration was 1mg/kg (diluted in saline 0,9% NaCl) and injected intraperitoneally. The control group was injected with a saline 0,9% NaCl solution at 10mL/kg.

### Viral tools

We used the different viruses present in the table. All viruses were diluted in phosphate-buffered saline (PBS) with 0.001% pluronic.

| Name of the virus | Company | Experiment |
| --- | --- | --- |
| AAV-PPTA-Cre | From Peter Vanhoutte | Fig. 2, 3, 4, 5, 6<br>Ext. Fig. 2, 6, 9 |
| AAV-PPE-Cre | From Peter Vanhoutte | Fig. 2, 3, 4, 5, 6<br>Ext. Fig. 2, 7, 9 |
| AAV5-hsyn-DIO-mCherry | Addgene | Fig. 1, 2, 3, 4, 5, 6<br>Ext. Fig. 1, 2, 3, 4, 5, 6, 7, 8, 9 |
| AAV5-EF1a-double floxed-hChR2(H134R)-mCherry-WPRE-HGHpA | Addgene | Fig. 3, 4, 5, 6<br>Ext. Fig. 4, 5, 9 |
| AAV8-hSyn-DIO-hM4D(Gi)-mCherry | Addgene | Fig. 7 |
| CAV-Cre | Plateforme de vectorologie de Montpellier | Fig. 1, 7<br>Ext. Fig. 1 |

### Stereotaxic injections

Stereotaxic injections were performed using a stereotaxic frame (Kopf Instruments) under general anesthesia with ketamine and xylazine (150mg/kg and 10mg/kg, respectively). All viruses were injected bilaterally at a rate of 100nL/min for a final volume of 500nL for dorsal hippocampus (dHPC) and 600nL for the nucleus accumbens (NAc) per site. Mice were 6 weeks-old (adult group) or 16 months-old (aged group) at the time of surgery and were given at least a 8-week recovery period to allow sufficient viral expression and reach the age of interest for this study, meaning 12 - 16 weeks old and 18 - 20 months old. Stereotaxic coordinates (antero-posterior: AP; mediolateral: ML; dorsoventral: DV) were chosen following^40^ and adapted regarding previous work of the team^56,78^. They are given here in millimetres (mm) from bregma for AP and ML coordinates, DV is taken from skull at the site of injection: dHPC: AP -2.1, ML ±1.7, DV -1.4 and NAc: AP +1.6, ML ±1.2, DV: -4.6.

To label the different neuronal populations of the NAc, notably D1 and D2 neurons, mice were injected with a mix of PPTA-Cre and AAV-DIO-mCherry or a mix of ENK-Cre and AAV-DIO-mCherry, respectively, in the NAc^44^. The PV-cre line was used to target the PV cells of the NAc, by injecting AAV-DIO-mCherry in this region. The projection from dHPC to NAc (dHPC^→NAc^) was studied with injection of AAV-ChR2-mCherry in the dHPC, allowing to express the channelrohdopsine in the CA1 neurons and enable the activation of the terminals by blue light pulses.

For projection-specific silencing (dHPC^→NAc^), wild-type mice were bilaterally injected with a retrograde CAV-2-Cre in the NAc and with a hSyn-DIO-hM4D(Gi)-mCherry in the dHPC (hereafter named dHPC^hM^^4→NAc^).

### Behavioral testing

In all behavioral tests, mice were housed in a habituation room adjacent to the experimental room at least one hour before the beginning of the test. In the experiment presented in figure 7i-k, Saline or CNO were administrated 30min before the CPP pre-test, with an IP injection. Mice were put back in the homecage box after the injection. All hippocampi were conserved and sliced in order to check the viral injections. If the injections were not located in the dorsal part of the hippocampus, the animals were excluded of the analysis.

#### O-maze

The O-maze test was used to measure anxiety levels. Each mouse was placed in a circular maze consisting of two open arms and two closed arms alternating in quadrants (width of walking lane: 5cm, total diameter: 55cm, height of the walls: 12cm, elevation above the floor: 60cm) with an ambient light of 100 lux in the open arms. Mice movements were recorded for 5 minutes using a video camera placed above the maze. The time spent in the open arms is scored to assess the level of anxiety.

#### Conditioned place preference (CPP)

The CPP protocol is adapted from Trouche et al., 2019^40^. The CPP apparatus consisted of two enclosures with distinct configurations: one square and one round (square: 46 cm × 46 cm × 38 cm). The two enclosures were connected via a bridge (8-cm length, 7-cm width) during CPP pre-test and test. The full maze was under an ambient light of 10 lux to avoid any anxious zone. All mice were handled for at least 3 days before the CPP experiment. Mice were housed by 2-3 per cage to avoid risk of fights and variability on the weight loss in-between mice due to hierarchy during food restriction. The weight was controlled every day at the same hour to maintain a ∼90% body weight and mice were identified by a non-invasive permanent marking on the tail. The two last days of restriction, they were exposed to few drops of 20% sucrose diluted in drinking water in a cup in the home cage to avoid novelty during the test. The fourth day, mice were submitted to the CPP. During pre-test, mice explored the entire apparatus for 10min, after which their spontaneous preference for one of the two enclosures was determined. After 15min back in their home cage, access to the bridge was removed for the conditioning sessions, and mice explored their ‘non-preferred’ enclosure in which we automatically delivered drops of 20% sucrose diluted in drinking water at a specific zone of the compartment. The duration of the conditioning was determined by the first consumption of sucrose, to which 10min were added, to ensure that the reward and compartment were associated. After another 15min break in their home cage, mice explored their ‘preferred’ enclosure in which drops of water were automatically delivered at a specific zone of the compartment, for the same time as the sucrose conditioning. One hour later (back in their home cage), the behavioural expression of the CPP memory was assessed by allowing the mice to explore the entire apparatus for 10min (CPP test). We calculated the place preference score for each mouse during both pre-test and test sessions as the difference between the time spent in the compartment paired with sucrose minus that paired with water during conditioning over their sum. The delta between the pre-test and the CPP test gave us the CPP score presented in the figures.

#### Sucrose preference

The sucrose preference test was performed in a separated stabling room under standard conditions, at 22°C, 55% to 65% humidity, with a 12-hour light/dark cycle (7am/7pm). The mice were isolated during the test from 6pm to 8am with food *ad libitum*, a plastic igloo, a chewing block of wood and cotton to facilitate nesting and avoid stress. Two bottles containing drinking water or 1% sucrose diluted in drinking water were at their disposal. The bottle configuration was different in between mice to counterbalance the preference of one side. Bottles were weighed before and after the test. The difference in weight indicates the amount of sucrose and water drunk, allowing us to calculate the percentage of sucrose preference.

### Patch-clamp electrophysiology

Electrophysiological recordings began at least one week after the last behavioural test. Whole-cell voltage-clamp configuration was used to measure synaptic responses, while current-clamp configuration was used to assess intrinsic properties of the cells such as intrinsic excitability, using an upright microscope (Olympus France). Patch-Clamp experiments were obtained using a Multiclamp 700B (Molecular Devices, Sunnyvale, CA). Signals were collected and stored using a Digidata 1550 B converter and pCLAMP 10.2 software (Molecular Devices, CA). After the whole-cell configuration obtention, neurons were left to stabilize for 2-3min before recordings began. Holding current and membrane resistance were monitored before and after the recordings, and if either of these two parameters varied by more than 20%, the recordings were discarded.

All hippocampi were conserved and sliced in order to check the viral injections. If the injections were not located in the dorsal part of the hippocampus, the recorded cells were excluded of the analysis.

#### Brain slice preparation

Mice were anesthetised (ketamine 150 mg/kg, xylazine 10 mg/kg) and transcardiacally perfused with ice-cold sucrose solution for slice preparation. Brains were cut on a vibratome (Microm HM600V, Thermo Scientific).

NAc slices: 250µm thick brain coronal slices were obtained in ice-cold dissecting solution containing (in mM): 234 sucrose, 2.5 KCl, 0.5 CaCl2, 10 MgCl2, 26 NaHCO3, 1.25 NaH2PO4 and 11 D-glucose, oxygenated with 95% O2 and 5% CO2, pH 7.4. Slices were first incubated, for 60 min at 37°C, in an artificial CSF (aCSF) solution containing (in mM): 119 NaCl, 2.5 KCl, 1.25 NaH2PO4, 26 NaHCO3, 1.3 MgSO4, 2.5 CaCl2 and 11 D-glucose, oxygenated with 95% O2 and 5% CO2, pH 7.4. Slices were used after recovering for another 30 min at room temperature. For all experiments, slices were perfused with the oxygenated aCSF at 31 ± 1°C. Hippocampal slices: 250 μm thick slices were obtained from previously isolated hippocampi and cut by using similar experimental conditions as for NAc slices.

#### Electrophysiology protocols

Recording pipettes (5-6 MΩ) for voltage-clamp experiments were filled with a solution containing the following (in mM): 117.5 Cs-gluconate, 15.5 CsCl, 10 TEACl, 8 NaCl, 10 HEPES, 0.25 EGTA, 4 MgATP and 0.3 NaGTP (pH 7.3; osmolarity 290-300 mOsm). For current-clamp experiments, the recording pipette solution contained (in mM): 135 gluconic acid (potassium salt: K-gluconate), 5 NaCl, 2 MgCl2, 10 HEPES, 0.5 EGTA, 2 MgATP and 0.4 NaGTP (pH 7.25; osmolarity 280-290 mOsm).

### Current-clamp

*Intrinsic excitability*: The resting membrane potential (Em) was first measured with I=0. Only cells with Em more negative than −55mV were considered. To study the relationship between firing frequency and current input, the neuron intrinsic excitability profile was accessed with Ih=0, i.e. at physiological resting membrane potential. Protocol: 200 ms duration with 50pA steps from -200pA to 500 pA. The membrane resistance was obtained by applying Ohm’s law equation to the potential obtained following -100pA current injection. The sag amplitude and sag kinetic were also defined with this negative current step taking the value of potential before the hyperpolarization and the peak value reached during the hyperpolarization. The firing rate (Hz) was calculated as the number of action potentials (APs) per second; the amplitude of the AP by measuring the delta between the beginning of the AP and its maximum peak for a 300pA injection.

### Voltage-Clamp

*Spontaneous activity*: for spontaneous excitatory post-synaptic currents (sEPSC) recordings, cells were clamped at -65mV and recorded for 5min. At least 100 events were obtained from each cell. Recordings were analysed with Clampfit 10.2 (Axon Instruments, U.S.A.) by applying a threshold search protocol to detecting spontaneous events, set at −8pA and 5ms duration excluding electric noise contamination in a software. We also measured spontaneous inhibitory post-synaptic currents (sIPSC). Cells were clamped at +10mV and recorded for 5min. Again, at least 100 events were obtained from each cell. Recordings were analysed by applying a threshold search to detecting spontaneous events, set at minimum +8pA amplitude and 5ms duration, thus excluding electric noise contamination. Amplitude and frequency of both sEPSC and sIPSC was calculated. Inhibitory and excitatory conductances were calculated by the following equations: [sEPSC amplitude x sEPSC frequency]/[Vh – Veq]; [sIPSC amplitude x sIPSC frequency]/[Vh – Veq], respectively. Vh=hold voltage; Veq=reversal potential of the current. The E/I ratio was obtained by the ratio between excitatory and inhibitory conductances.

*Optogenetics:* To assess the dHPC to NAc projection (dHPC^→NAc^), we used electrophysiology couple to optogenetic. We used viral tools to label the different cell populations of the NAc while opto-genetically activating the dHPC terminals. 470nm light pulses from a LED source (CoolLED pE-100) were delivered directly through the microscope’s objective. Optically evoked excitatory post-synaptic currents (oEPSC) were obtained every 20s with 8mW and 5ms light pulses, while the cell was clamped at -65mV. The amplitude and latency of the response was calculated with the average of 20 consecutive traces. The optically evoked inhibitory post-synaptic current (oIPSC) were obtained with the same protocol but the cell was clamped at +10mV. To assess the neurotransmitter nature of the evoked response, we used the same protocol described for oEPSC and oIPSC and pharmacologically isolated the glutamatergic component using picrotoxin (50µM) to block GABAa receptors or isolated GABAergic component of the optogenetic stimulation using AMPA receptor antagonists (DNQX 10 µM).

In all cases, offline analysis was performed using Clampfit 10.2 (Axon Instruments, U.S.A.).

### Histology

#### Brain slice preparation

Both adult and old mice were anesthetised (ketamine 150 mg/kg, xylazine 10 mg/kg) and transcardiacally perfused with PFA solution for tissue fixation. Brains were cut into 15-μm coronal sections on a vibratome (Microm HM650V, Thermo Scientific).

#### Viral injection confirmation

After electrophysiological recordings, hippocampi were cut into 50µm coronal section on a vibratome. Using an epifluorescence microscope, dHPC was imaged to verify the viral injections. If the injections were not located in the CA1 part of the hippocampus, the recorded cells were excluded of the analysis.

#### Immunofluorescence

Assays were performed in triplicate on both hemispheres of the HPC and NAc (n = 8 per group). Sections were washed tree times for 5 min in PBS 1x before being blocked for 30 minutes in staining solution (PBS 1x, 0.3% Tritton X-100 in, 0.5% BSA and 10% Horse Serum) at room temperature. Then, sections were incubated with primary antibodies in staining solution overnight at 4°C with agitation; Sections were then washed tree times for 5 min in PBS 1x before and after being incubated with secondary antibodies in staining solution for 2 hours at room temperature. After PBS 1x washes, 4,6-diamidino-2-phenylindole (DAPI, 1:20000; Roche, 10236276001) was used for nuclei staining for 5 min at room temperature. Sections were washed with PSB 1x and mounted with Mowiol (Sigma 81381) and stored at 4°C until imaging. Images were taken using a 20x objective at EVOS (Invitrogen M7000) microscope and Videomicroscope (Zeiss) and DsRed staining was manually analyzed, whereas PV and DAPI analyses were performed by QuPath (v.0.5.1).

| Name of the antibody | Company | Identifier |
| --- | --- | --- |
| Mouse monoclonal anti-Parvalbumin (1:1000) | Sigma | Cat#P3088 |
| Rabbit polyclonal anti-DsRed (1:2000) | Clontech | Cat#632496 |
| Goat anti-mouse IgG - Alexa Fluor 488 (1:1000) | Life tech | Cat#A11029 |
| Donkey anti-rabbit IgG - Alexa Fluor 594 (1:1000) | Invitrogen | Cat#A21207 |

### Random Forest and Principal Component Analysis

#### Data Collection and Preprocessing

The dataset comprised behavioral and electrophysiological measurements from adult (3-4 months, n=13) and aged (18-20 months, n=12) C57BL/6 mice. Variables included behavioral scores (sucrose preference, CPP score for compartment and exploration), electrophysiological parameters (oEPSC amplitude, oIPSC amplitude, sEPSC frequency, sIPSC frequency, paired-pulse ratio), connectivity ratio, and synaptic fatigue measurements, totalling 13 features initially. Missing data were addressed through mean imputation for variables with <5% missing values. Variables with excessive missing data (>20%), including O-maze scores, social interaction scores, and certain synaptic current measurements, were excluded from subsequent analyses to maintain data integrity. All continuous variables were standardized (z-score normalization) prior to multivariate analysis to account for differences in measurement scales.

#### Principal Component Analysis

Principal Component Analysis (PCA) was performed using Python’s scikit-learn library (v1.3.0) to reduce dimensionality and identify patterns distinguishing adult from aged mice. PCA was initially conducted on the combined dataset including both D1R-MSN and D2R-MSN subtypes. Following preliminary analysis revealing clearer age-related separation in D1R-MSN, subsequent PCA focused exclusively on the D1R-MSN subset with 10 retained features. The analysis decomposed the standardized data into orthogonal components representing maximum variance. Results were visualized using biplots displaying both sample scores and variable loadings on the first two principal components (PC1 and PC2). The contribution of each original variable to the principal components was assessed through loading vectors. Multivariate analysis of variance (MANOVA) was applied to test statistical significance of group separation (α = 0.05).

#### Random Forest Classification

A Random Forest classifier was implemented using scikit-learn’s RandomForestClassifier to identify variables most predictive of age group classification. The dataset was partitioned into training (70%) and testing (30%) subsets using stratified random sampling to maintain proportional representation of age groups. The Random Forest model consisted of 100 decision trees constructed through bootstrap aggregation. Each tree was built using a random subset of features at each split, with all other hyperparameters set to default values. Model performance was evaluated on the held-out test set using accuracy metrics. Feature importance was quantified using mean decrease in Gini impurity, which measures each variable’s contribution to classification accuracy across all decision trees. Variables were ranked by importance scores, and Mann-Whitney U tests were performed post-hoc to validate differences in top-ranked features between age groups (α = 0.05). Comparative analysis was conducted between Random Forest performance on original scaled data versus PCA-transformed data to assess information retention through dimensionality reduction.

## Data analysis

Data were analysed using Prism (GraphPad, U.S.A.). Outliers were identified by applying the ROUT method (Q=1%) by using the defined software. Normality of the distribution was first tested using the four tests: Anderson-Darling test, D’Agostino and Pearson test, Shapiro-Wilk test and Kolmogorov-Smirnov test. The data were considered non-normally distributed if two or more measures failed to meet normality criteria in any group. Depending of the number of groups and the results of the normality test, groups were then compared using a Student t-test, U of Mann-Whitney, or two-way analyses of variance (ANOVA) followed by post hoc Tuckey test. For proportion analysis, Chi-square or Fisher-test were used. Statistical significance was achieved as p<0.05. Data in the figures are presented as mean ± SEM. N= number of animals. n = number of cells.

## Author contributions

AR, JB and PAP. designed the study and interpreted the results. A.R. performed all *in vivo* stereotaxic surgeries, electrophysiological recordings, behavioral assays assisted by TB and IB and data analysis, except when otherwise stated. BFD performed immunofluorescence assays assisted by MV. LY made fMRI analysis assisted by SR. ED performed Principal Component and Random Forest Analysis. MDS made patch-clamp electrophysiological recordings to investigate the E/I ratio in the hippocampus. PV contributed with viral tools and help on experimental design, as also LVL. PAP coordinated the project. AR wrote the manuscript with input from the other authors. All authors have revised the manuscript, discussed the experimental findings, and approved the final version.

Funding and acknowledgments:

This work was supported by the Agence National de Recherche (ANR-22-CE37-0017-01). AR is a recipient of a PhD fellowship from the Ministère de la Recherche, de l’Enseignement Supérieur et de l’Innovation and the French government through the France 2030 investment plan managed by the National Research Agency (ANR), as part of the Initiative of Excellence Université Côte d’Azur under reference number ANR- 15-IDEX-01. ED was funded by the interdisciplinary Institute for Modeling in Neuroscience and Cognition (NeuroMod) of the University Côte d’Azur (MSc Mod4NeuCog). LVL and MDS are supported by Fundação para a Ciência e Tecnologia, Portugal (UI/BD/154567/2022).

---

**Extended data Figure 1.**
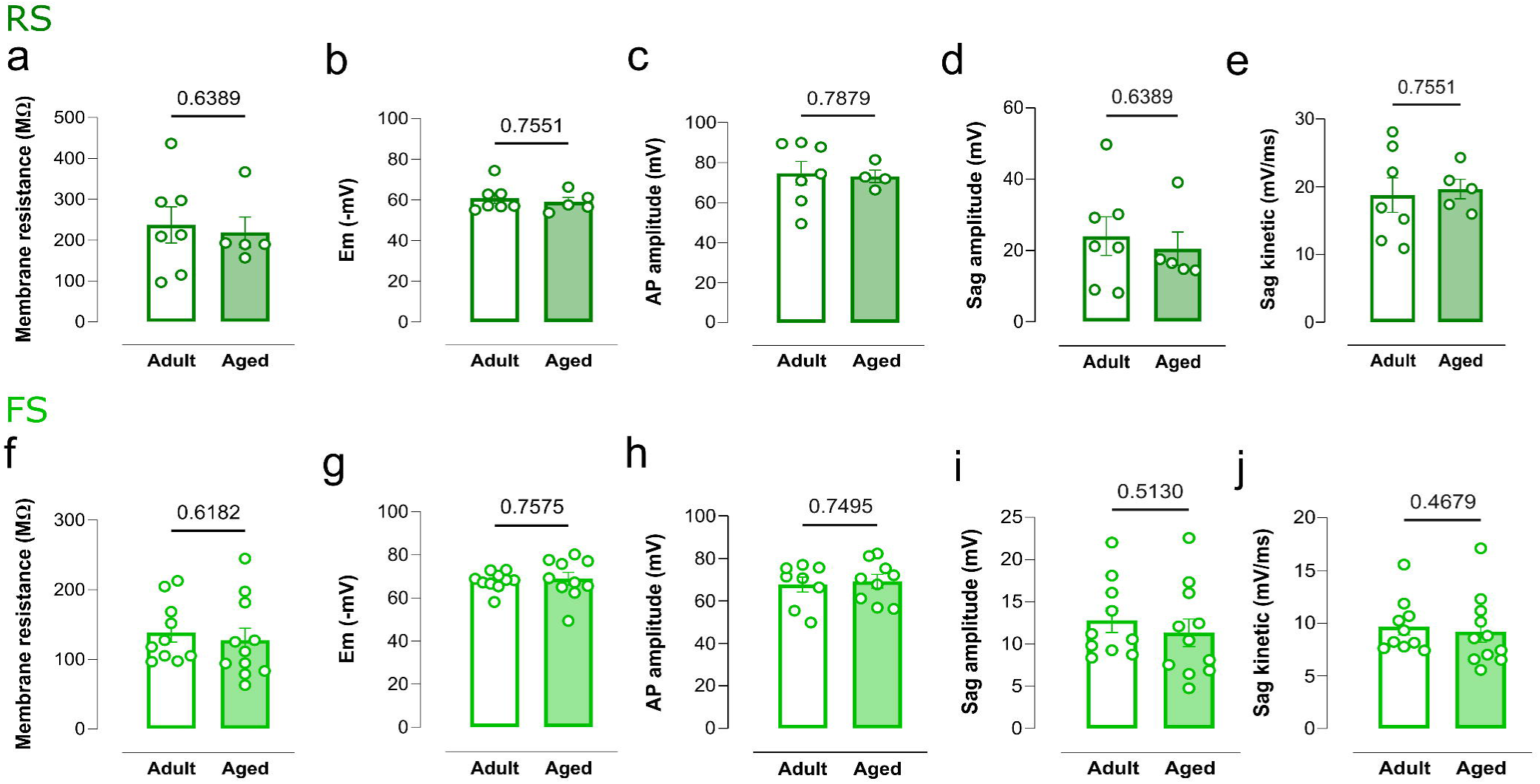
: Intrinsic properties of both regular- (RS) and fast-spiking (FS) projecting cells are preserved in aging. (a) Membrane resistance shows no differences. Results are expressed as the mean ± SEM. (Normality test not passed. Mann-Whitney test, p = 0.6389, adult N= 5, n = 7, aged N= 4, n = 5). **(b)** Membrane potential (Em) was similar between adult and aged mice. Results are expressed as the mean ± SEM. (Normality test not passed. Mann-Whitney test, p = 0.7551, adult N= 5, n = 7, aged N= 4, n = 5). **(c)** Action potential (AP) amplitude showed no differences in aging for regular spiking projecting cells. Results are expressed as the mean ± SEM. (Normality test not passed. Mann-Whitney test, p = 0.7879, adult N= 5, n = 7, aged N= 3, n = 4). **(d)** Sag amplitude was similar between adult and aged. Results are expressed as the mean ± SEM. (Normality test not passed. Mann-Whitney test, p = 0.6389, adult N= 5, n = 7, aged N= 3, n = 5). **(e)** Sag kinetic showed no differences in aging. Results are expressed as the mean ± SEM. (Normality test not passed. Mann-Whitney test, p = 0.7551, adult N= 5, n = 7, aged N= 3, n = 5). **(f-j)** Fast-spiking (FS) cells show similar pattern as regular-spiking cells as no differences were observed. **(f)** Results are expressed as the mean ± SEM. (Passed normality test. Unpaired t-test, p = 0.6182, adult N= 7, n = 10, aged N= 7, n = 11). **(g)** Results are expressed as the mean ± SEM. (Passed normality test. Unpaired t-test, p = 0.7575, adult N= 7, n = 10, aged N= 7, n = 11). **(h)** Results are expressed as the mean ± SEM. (Passed normality test. Unpaired t-test, p = 0.7495, adult N= 7, n = 8, aged N= 7, n = 9). **(i)** Results are expressed as the mean ± SEM. (Passed normality test. Unpaired t-test, p = 0.5130, adult N= 7, n = 10, aged N= 7, n = 11). **(n)** Results are expressed as the mean ± SEM. (Normality test not passed. Mann-Whitney test, p = 0.4679, adult N= 7, n = 10, aged N= 7, n = 11). In all panels: significant p-values are represented in bold, N = number of animals, n = number of cells.

**Extended data Figure 2.**
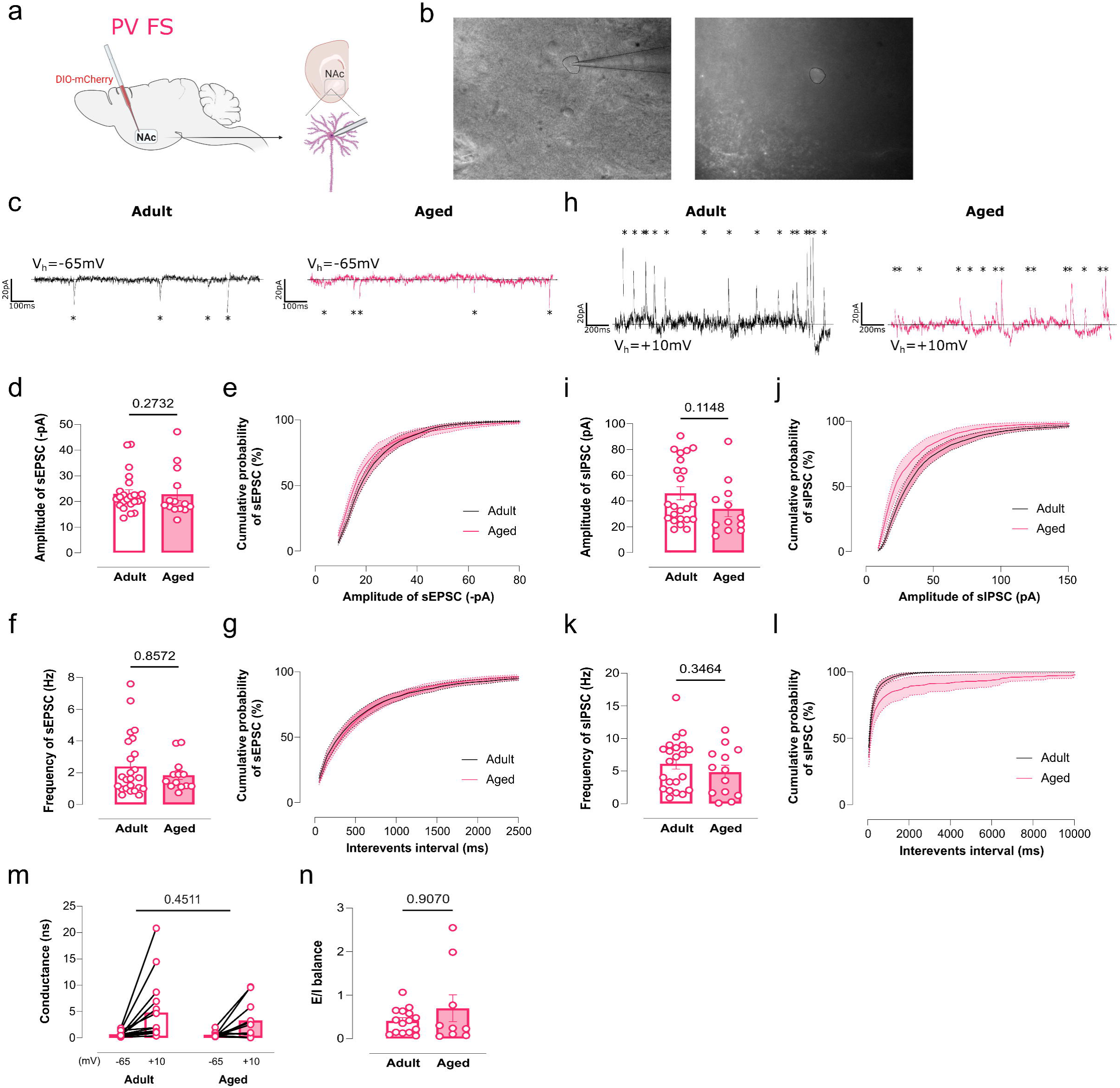
: Spontaneous currents cumulative histograms. The cumulative probability of recorded currents shown in Fig.2 follows for D1R-MSN: (a) sEPSC amplitude in D1R-MSN, (b) sEPSC frequency in D1R-MSN, (c) sIPSC amplitude in D1R-MSN, (d) sIPSC frequency in D1R-MSN. (e) Synaptic conductances of D1R-MSN, derived from currents recorded at holding potentials of −65 mV for excitatory and +10 mV for inhibitory inputs were maintained upon aging. Results are expressed as the mean ± SEM. (Two-way ANOVA, interaction: p = 0.8773, F(1, 50) = 0.02407, adult N = 9, n = 28, aged N = 8, n = 24). (f) The E/I ratio is preserved in aging. Results are expressed as the mean ± SEM. (Passed normality test. Unpaired t-test, p = 0.7922, adult N = 9, n = 26, aged N = 8, n = 20). The cumulative probability of recorded currents shown in Fig.2 follows for D2R-MSN: (g) sEPSC amplitude in D2R-MSN, (h) sEPSC frequency in D2R-MSN, (i) sIPSC amplitude in D2R-MSN, (j) sIPSC frequency in D2R-MSN. (k) Synaptic conductances of D2R-MSN, derived from currents recorded at holding potentials of −65 mV for excitatory and +10 mV for inhibitory inputs were maintained upon aging. Results are expressed as the mean ± SEM. (Two-way ANOVA, interaction: p = 0.7787, F(1, 36) = 0.08017, adult N = 9, n = 10, aged N = 8, n = 28). (l) The E/I ratio is maintained in aging for D2R-MSN. Results are expressed as the mean ± SEM. (Normality test not passed. Mann-Whitney test, p = 0.1014, adult N = 9, n = 17, aged N = 8, n = 27).

**Extended data Figure 3.**
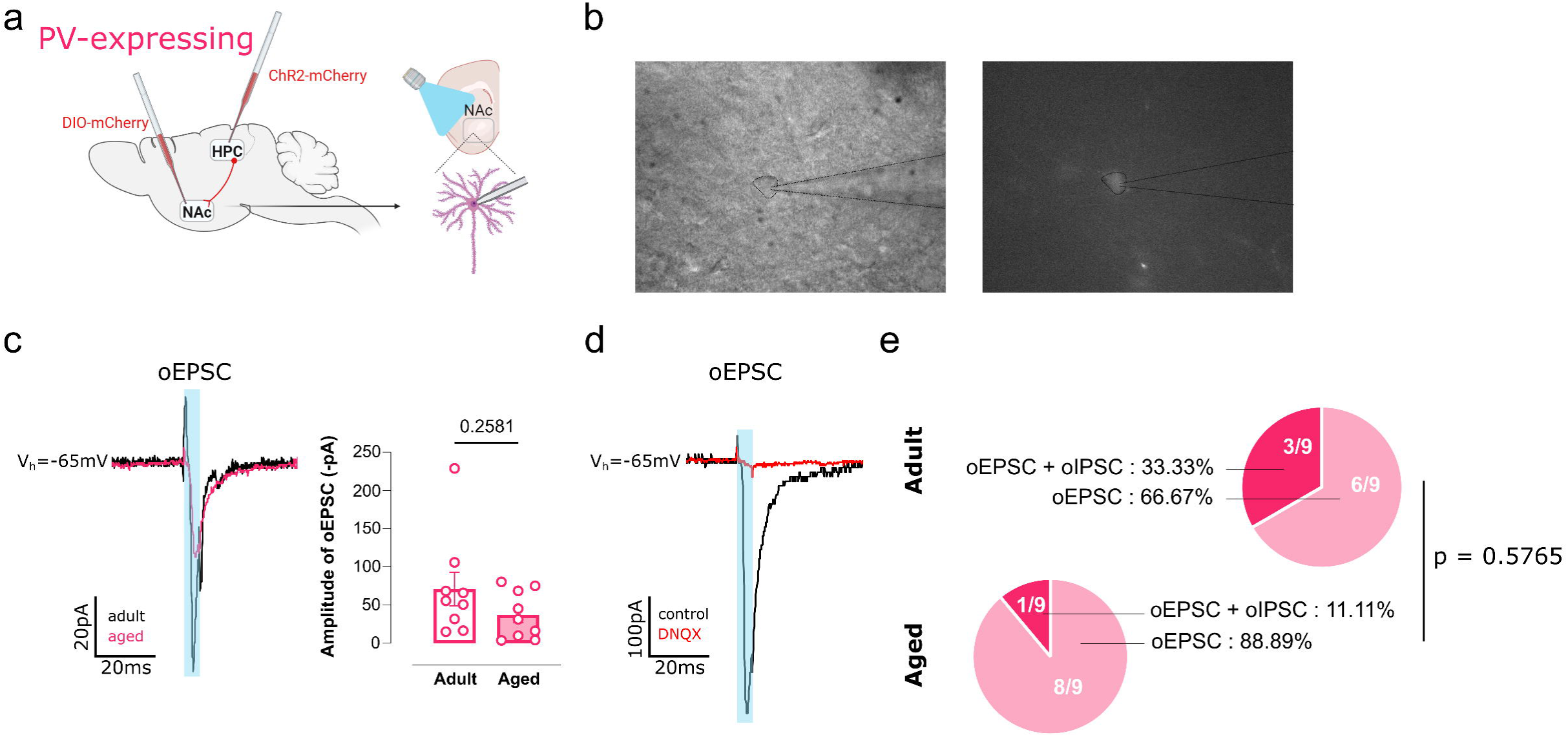
: PV-expressing neurons of the NAc are preserved with aging. **(a)** Schematic representation of stereotaxic injections: viral tracing with DIO-mCherry injected in the NAc of PV-Cre mice to label specifically the PV-expressing population. **(b)** Representative images of the PV-expressing neurons with a recorded pipette, without and with fluorescence. **(c)** Representative traces of spontaneous excitatory post synaptic currents (sEPSC) for adult and aged PV-expressing neurons. **(d)** Amplitude of sEPSC is similar in both groups. Results are expressed as the mean ± SEM. (Normality test not passed. Mann-Whitney test, p = 0.2732, adult N = 7, n = 24, aged N = 4, n = 14). **(e)** Cumulative distribution of sEPSC amplitude in PV-expressing neurons. **(f)** Frequency of sEPSC is increased in aged PV-expressing neurons. Results are expressed as the mean ± SEM. (Normality test not passed. Mann-Whitney test, p = 0.8572, adult N = 7, n = 24, aged N = 4, n = 13). **(g)** Cumulative distribution of sEPSC frequency. **(h)** Representative traces of spontaneous inhibitory post synaptic currents (sIPSC) for adult and aged PV-expressing neurons. **(i)** Amplitude of sIPSC is preserved for aged PV-expressing neurons. Results are expressed as the mean ± SEM. (Normality test not passed. Mann-Whitney test, p = 0.1148, adult N = 7, n = 23, aged N = 4, n = 12). **(j)** Cumulative distribution of sIPSC amplitude in PV-expressing neurons. **(k)** Frequency of sIPSC is maintained in aging. Results are expressed as the mean ± SEM. (Passed normality test. Unpaired t-test, p = 0.3464, adult N = 7, n = 22, aged N = 4, n = 13). **(l)** Cumulative distribution of sIPSC frequency in PV-expressing neurons. **(m)** Synaptic conductances of PV-expressing neurons were derived from currents recorded at holding potentials of −65 mV for excitatory and +10 mV for inhibitory inputs. Results are expressed as the mean ± SEM. (Two-way ANOVA, p = 0.4511, F(1, 24) = 0.5869, adult N = 7, n = 15, aged N = 4, n = 11). **(n)** The E/I ratio is maintained in aging. Results are expressed as the mean ± SEM. (Normality test not passed. Mann-Whitney test. p = 0.9070, adult N = 7, n = 15, aged N = 4, n = 9). In all panels: significant p-values are represented in bold, N = number of animals, n = number of cells.

**Extended data Figure 4.**
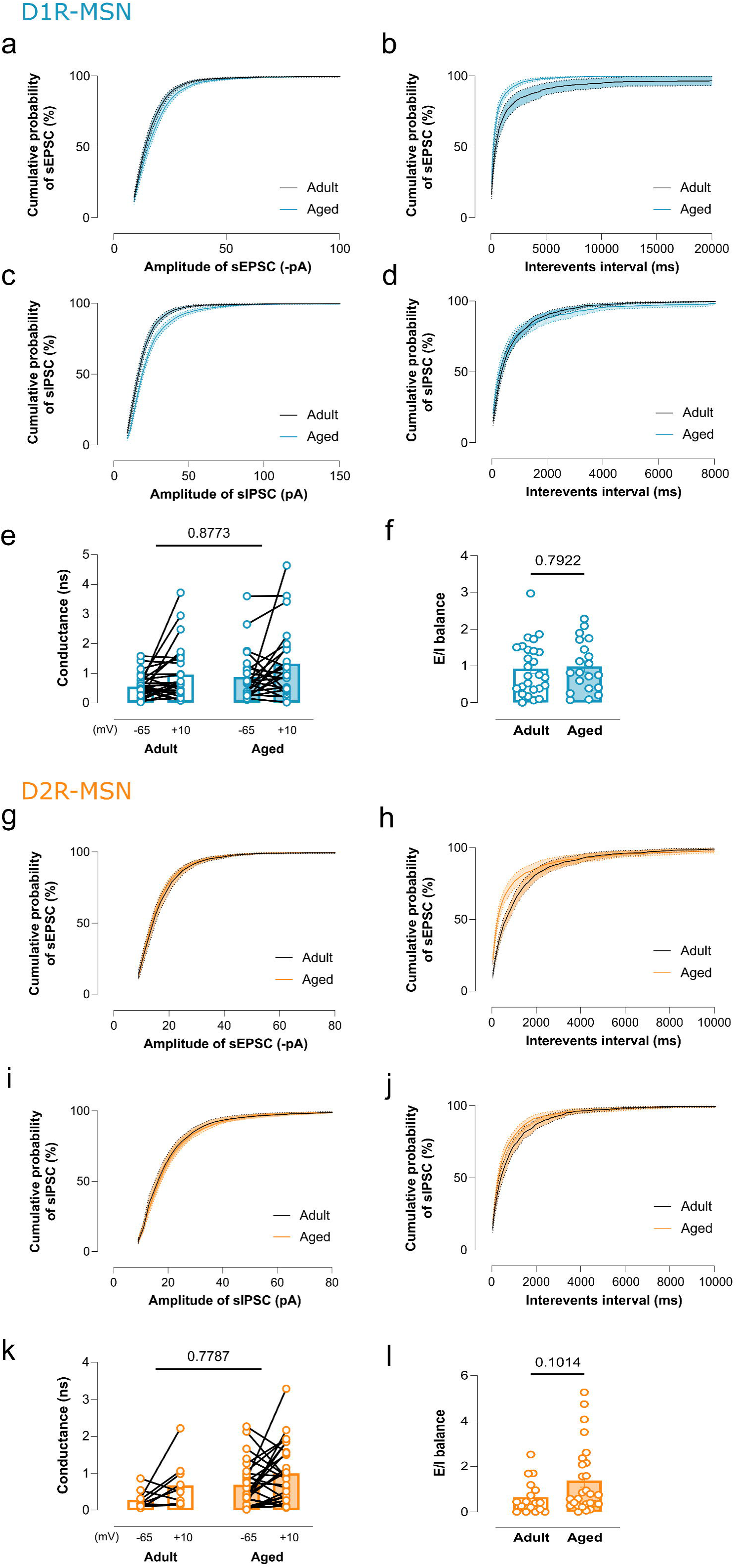
: dHPC^◊PV-expressing^ ^neurons^ pathway is preserved in aged males. (a) Schematic representation of stereotaxic injections: viral tracing with DIO-mCherry injected in the NAc of PV-Cre mice to label specifically the PV-expressing population and AAV-ChR2-mCherry in the dHPC. **(b)** Representative images of a PV-expressing cell with a recording pipette, without and with fluorescence. **(c)** Representative traces of optogenetically-evoked excitatory post synaptic currents (oEPSC) after a 5ms blue light stimulation. Amplitude of oEPSC is maintained in aged PV-expressing neurons. Results are expressed as the mean ± SEM. (Normality test not passed. Mann-Whitney test, p = 0.2581, adult N = 4, n = 9, aged N = 4, n = 9). **(d)** Representative traces of optogenetic stimulation of the dHPC terminals eliciting oEPSC in PV-expressing cells before (black) and after DNQX (red), a glutamatergic antagonist revealing the glutamatergic nature of the dHPC^→PV-expressing neurons^projection. **(e)** The proportion of PV-expressing neurons eliciting oEPSC or both oEPSC and oIPSC is maintained in aging. Results are expressed as percentages. (Fischer’s exact test, p = 0.5765, adult N = 4, n = 9, aged N = 4, n = 9). In all panels: significant p-values are represented in bold, N = number of animals, n = number of cells.

**Extended data Figure 5.**
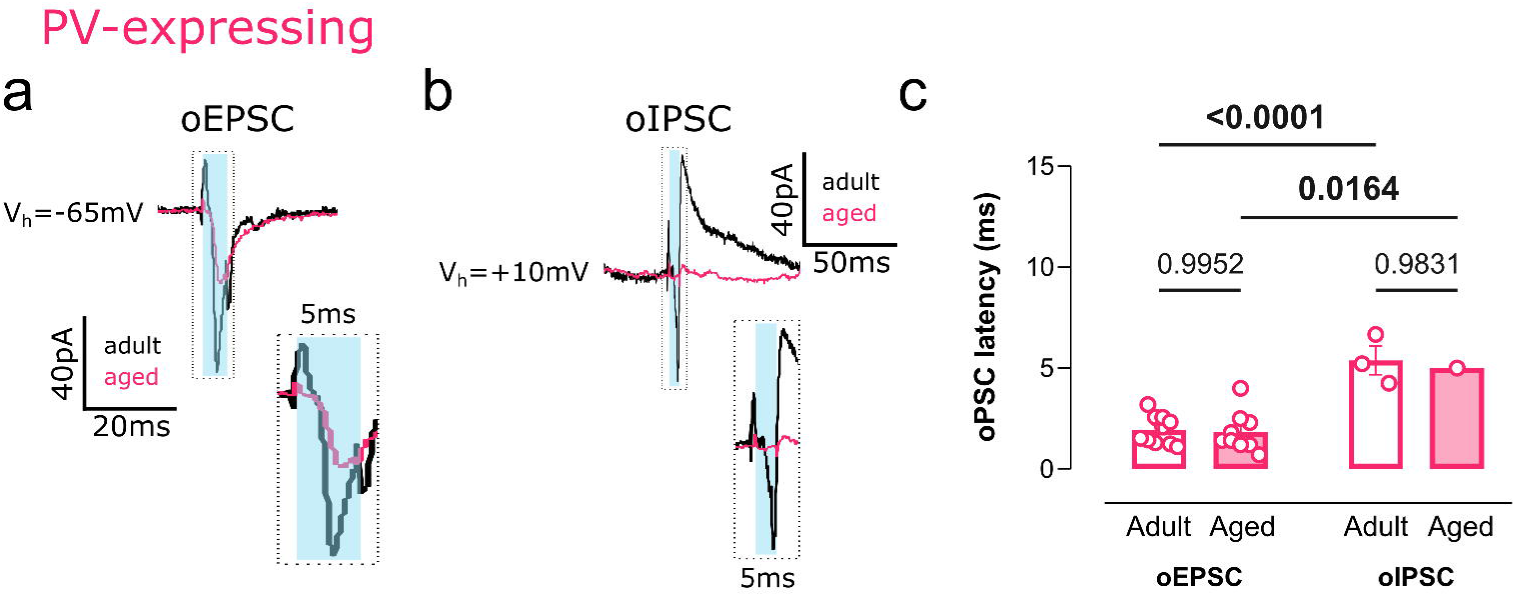
: Inhibitory responses of PV-expressing neurons of the NAc also exhibit increased latency. **(a)** Representative traces of optogenetically-evoked excitatory post synaptic currents (oEPSC) after a 5ms blue light stimulation, represented as blue bar. Right: enlarged illustration of the dashed rectangle. Blue bar represents the optical stimulation. **(b)** Representative traces of optogenetically-evoked inhibitory post synaptic currents (oIPSC) after a 5ms blue light stimulation, represented as blue bar. Right: enlarged illustration of the dashed rectangle. Blue bar represents the optical stimulation. **(c)** Latency of oEPSC and oIPSC is maintained during aging for PV-expressing neurons, but oIPSC show a higher latency (around 5ms) compared to oEPSC (around 2ms). Results are expressed as the mean ± SEM. (Two-way ANOVA, V hold: p < 0.0001, F (1, 19) = 35,08, oEPSC adult N = 4, n = 10, aged N = 3, n = 9; oIPSC adult N = 1, n = 3, aged N = 1, n = 1). In all panels: significant p-values are represented in bold, N = number of animals, n = number of cells.

**Extended data Figure 6.**
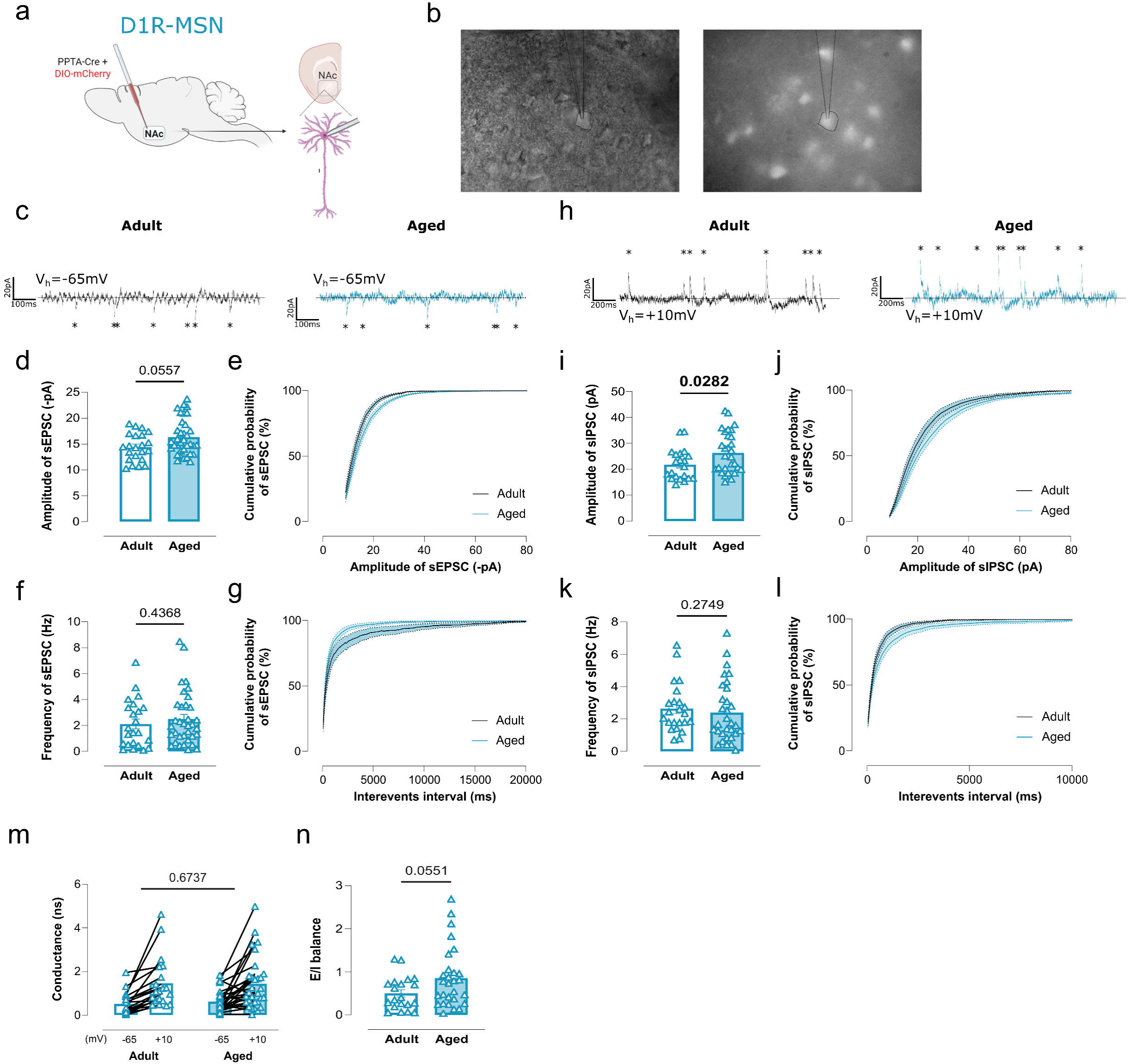
: Inhibitory spontaneous transmission is increased for D1R-MSN **in aged female.** (a) Schematic representation of stereotaxic injections: viral tracing with PPTA-Cre combined with DIO-mCherry injected in the NAc to label specifically D1R-MSN population. **(b)** Representative images of the D1R-MSN with a recorded pipette, without and with fluorescence. **(c)** Representative traces of spontaneous excitatory post synaptic currents (sEPSC) for adult and aged D1R-MSN. **(d)** Amplitude of sEPSC is similar in both groups. Results are expressed as the mean ± SEM. (Normality test not passed. Mann-Whitney test, p = 0.0557, adult N = 7, n = 22, aged N = 8, n = 35). **(e)** Cumulative distribution of sEPSC amplitude in D1R-MSN. **(f)** Frequency of sEPSC is maintained for D1R-MSN in females. Results are expressed as the mean ± SEM. (Normality test not passed. Mann-Whitney test, p = 0.4368, adult N = 7, n = 23, aged N = 8, n = 35). **(g)** Cumulative distribution of sEPSC frequency in D1R-MSN. **(h)** Representative traces of spontaneous inhibitory post synaptic currents (sIPSC) for adult and aged D1R-MSN in females. **(i)** Amplitude of sIPSC is increased for aged D1R-MSN. Results are expressed as the mean ± SEM. Passed normality test. Unpaired t-test, p = 0.0282, adult N = 7, n = 21, aged N = 8, n = 31). **(j)** Cumulative distribution of sIPSC amplitude in D1R-MSN. **(k)** Frequency of sIPSC is maintained with aging. Results are expressed as the mean ± SEM. (Normality test not passed. Mann-Whitney test, p = 0.2749, adult N = 7, n = 23, aged N = 8, n = 31). **(l)** Cumulative distribution of sIPSC frequency in D1R-MSN. **(m)** Synaptic conductances of D1R-MSN in females were derived from currents recorded at holding potentials of −65 mV for excitatory and +10 mV for inhibitory inputs. Results are expressed as the mean ± SEM. (Two-way ANOVA, p = 0.6737, F(1, 48) = 0.1795, adult N = 7, n = 20, aged N = 8, n = 30). **(n)** The E/I ratio is maintained in aging. Results are expressed as the mean ± SEM. (Normality test not passed. Mann-Whitney test, p = 0.0551, adult N = 7, n = 20, aged N = 8, n = 28). In all panels: significant p-values are represented in bold, N = number of animals, n = number of cells.

**Extended data Figure 7.**
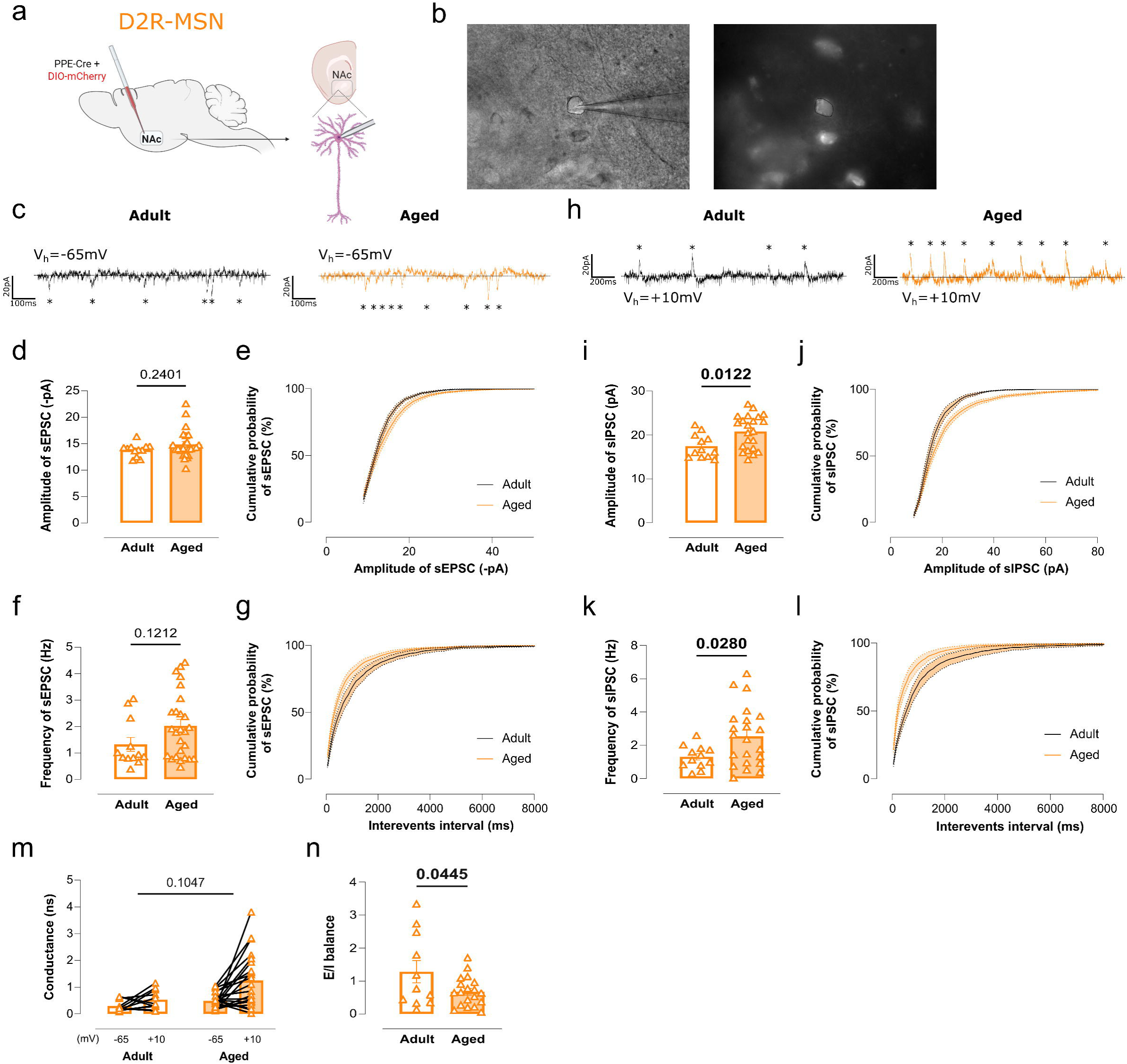
: Inhibitory spontaneous transmission is increased in D2R-MSN of aged female. **(a)** Schematic representation of stereotaxic injections: viral tracing with PPE-Cre combined with DIO-mCherry injected in the NAc to label specifically D2R-MSN population. **(b)** Representative images of the D2R-MSN with a recorded pipette, without and with fluorescence. **(c)** Representative traces of spontaneous excitatory post synaptic currents (sEPSC) for adult and aged D2R-MSN. **(d)** Amplitude of sEPSC is similar in both groups. Results are expressed as the mean ± SEM. (Normality test not passed. Mann-Whitney test, p = 0.2401, adult N = 3, n = 12 aged N = 7, n = 25). **(e)** Cumulative distribution of sEPSC amplitude in D2R-MSN. **(f)** Frequency of sEPSC is maintained for D2R-MSN in females. Results are expressed as the mean ± SEM. (Normality test not passed. Mann-Whitney test, p = 0.1212, adult N = 3, n = 12, aged N = 7, n = 26). **(g)** Cumulative distribution of sEPSC frequency in D2R-MSN. **(h)** Representative traces of spontaneous inhibitory post synaptic currents (sIPSC) for adult and aged D2R-MSN in females. **(i)** Amplitude of sIPSC is increased for aged D2R-MSN. Results are expressed as the mean ± SEM. (Passed normality test. Unpaired t-test, p = 0.0122, adult N = 3, n = 12, aged N = 7, n = 22). **(j)** Cumulative distribution of sIPSC amplitude in D2R-MSN. **(k)** Frequency of sIPSC is increased with aging. Results are expressed as the mean ± SEM. (Passed normality test. Unpaired t-test, p = 0.0280, adult N = 3, n = 12, aged N = 7, n = 22). **(l)** Cumulative distribution of sIPSC frequency in D2R-MSN. **(m)** Synaptic conductances of D2R-MSn in females were derived from currents recorded at holding potentials of −65 mV for excitatory and +10 mV for inhibitory inputs. Results are expressed as the mean ± SEM. (Two-way ANOVA, p = 0.1047, F(1, 31) = 2.794, adult N = 3, n = 11, aged N = 7, n = 22). **(n)** The E/I ratio is decreased in aging, towards greater inhibition. Results are expressed as the mean ± SEM. (Passed normality test. Unpaired t-test, p = 0.0445, adult N = 3, n = 11, aged N = 7, n = 19). In all panels: significant p-values are represented in bold, N = number of animals, n = number of cells.

**Extended data Figure 8.**
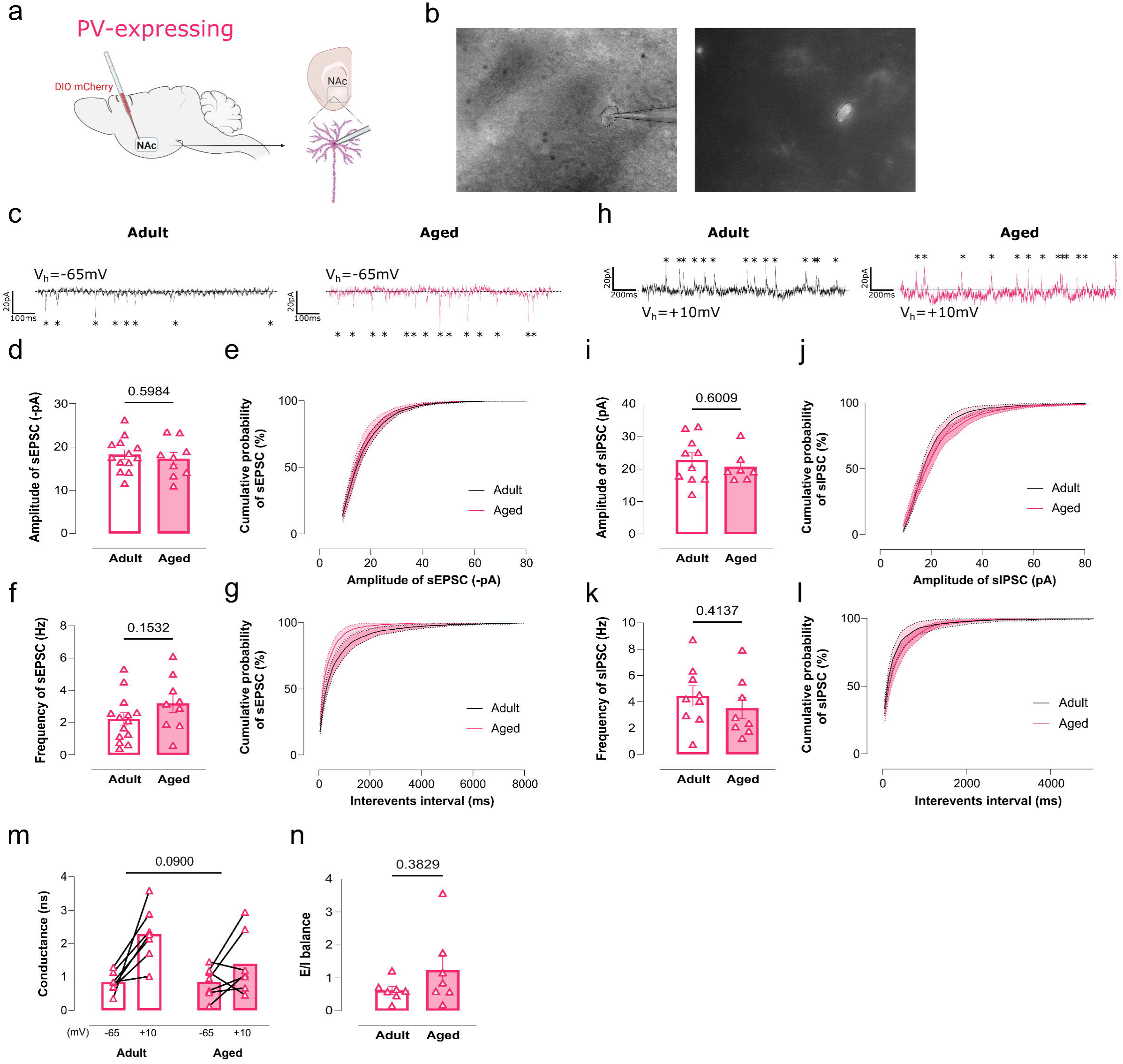
: Spontaneous transmission is maintained in PV-expressing neurons of aged female. **(a)** Schematic representation of stereotaxic injections: viral tracing with DIO-mCherry injected in the NAc of PV-Cre mice to label specifically PV-expressing population. **(b)** Representative images of the PV-expressing neuron with a recorded pipette, without and with fluorescence. **(c)** Representative traces of spontaneous excitatory post synaptic currents (sEPSC) for adult and aged PV-expressing neuron. **(d)** Amplitude of sEPSC is similar in both groups. Results are expressed as the mean ± SEM. (Passed normality test. Unpaired t-test, p = 0.5984, adult N = 5, n = 13 aged N = 4, n = 9). **(e)** Cumulative distribution of sEPSC amplitude in PV-expressing neurons. **(f)** Frequency of sEPSC is maintained for PV-expressing neurons in females. Results are expressed as the mean ± SEM. (Passed normality test. Unpaired t-test, p = 0.1532, adult N = 5, n = 14, aged N = 4, n = 9). **(g)** Cumulative distribution of sEPSC frequency in PV-expressing neurons. **(h)** Representative traces of spontaneous inhibitory post synaptic currents (sIPSC) for adult and aged PV-expressing neurons in females. **(i)** Amplitude of sIPSC is maintained for aged PV-expressing neurons. Results are expressed as the mean ± SEM. (Normality test not passed. Mann-Whitney test, p = 0.6009, adult N = 5, n = 10, aged N = 4, n = 7). **(j)** Cumulative distribution of sIPSC amplitude in PV-expressing neurons. **(k)** Frequency of sIPSC is preserved with aging. Results are expressed as the mean ± SEM. (Passed normality test. Unpaired t-test, p = 0.4137, adult N = 5, n = 9, aged N = 4, n = 8). **(l)** Cumulative distribution of sIPSC frequency in PV-expressing neurons. **(m)** Synaptic conductances of PV-expressing neurons of females were derived from currents recorded at holding potentials of −65 mV for excitatory and +10 mV for inhibitory inputs. Results are expressed as the mean ± SEM. (Two-way ANOVA, p = 0.0900, F(1, 12) = 3.401, adult N = 5, n = 7, aged N = 4, n = 7). **(n)** The E/I ratio is maintained in aging. Results are expressed as the mean ± SEM. (Normality test not passed. Mann-Whitney test, p = 0.3829, adult N = 5, n = 7, aged N = 4, n = 7). In all panels: significant p-values are represented in bold, N = number of animals, n = number of cells.

**Extended data Figure 9.**
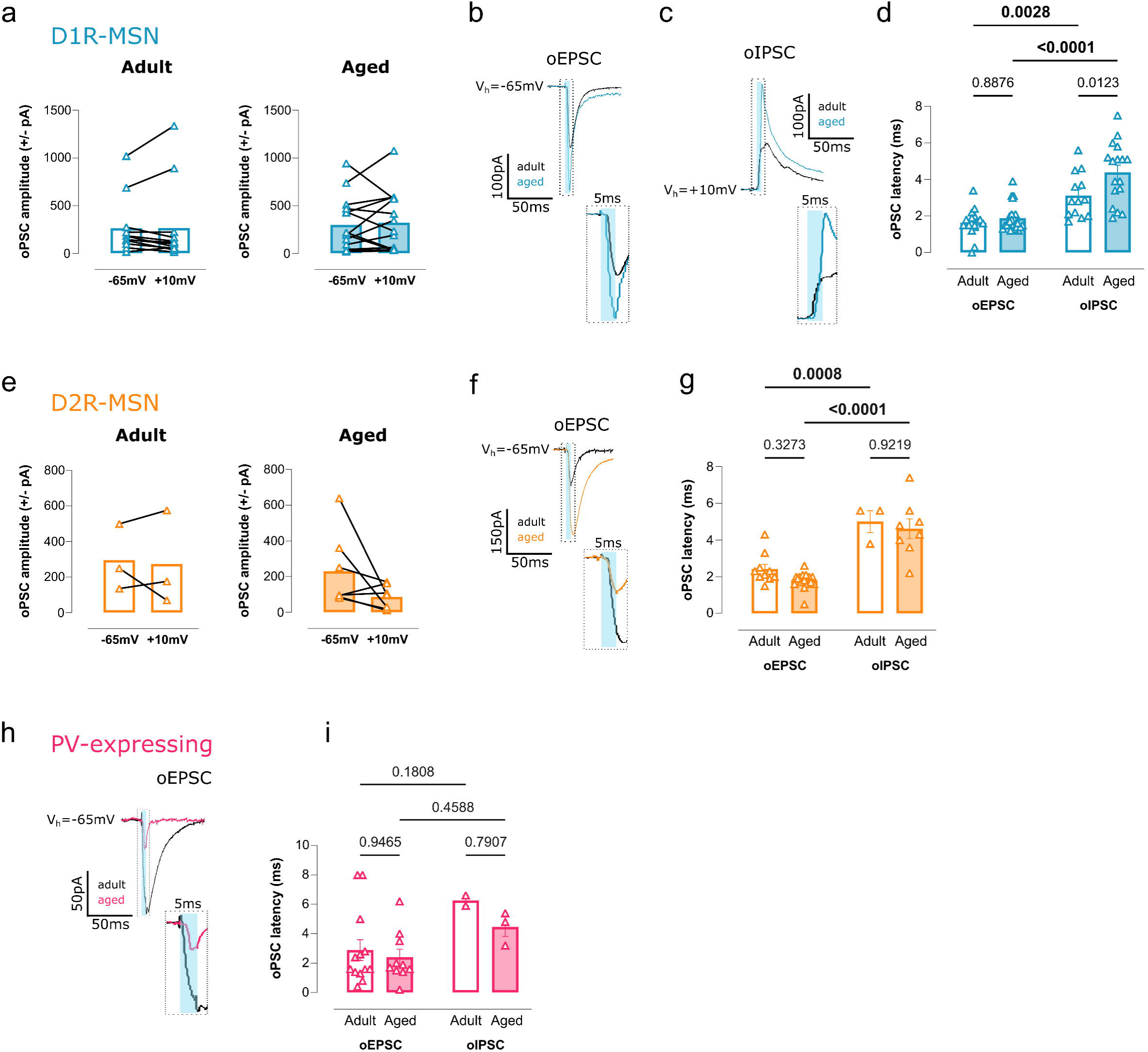
: The latency in the inhibitory response is conserved in females in the different cell types. **(a)** Optogenetically-evoked post synaptic currents for the same cells at V_h_ = -65mV (excitatory) and V_h_ = -10mV (inhibitory). Results are expressed as the mean ± SEM. (For adult group: N = 5, n = 12. For aged group: N = 9, n =15). **(b)** Representative traces of optogenetically-evoked excitatory post synaptic currents (oEPSC) after a 5ms blue light stimulation, represented as blue bar. Right: enlarged illustration of the dashed rectangle. Blue bar represents the optical stimulation. **(c)** Representative traces of optogenetically-evoked inhibitory post synaptic currents (oIPSC) after a 5ms blue light stimulation, represented as blue bar. Right: enlarged illustration of the dashed rectangle. Blue bar represents the optical stimulation. **(d)** Latency of oEPSC and oIPSC is maintained during aging for D1R-MSN in females, but oIPSC show a higher latency (around 5ms) compared to oEPSC (around 2ms). Results are expressed as the mean ± SEM. Results are expressed as the mean ± SEM. (Two-way ANOVA, age: p = 0.054, F(1, 68) = 8.260; V hold: p < 0.0001, F(1, 68) = 56.12, oEPSC adult N = 8, n = 17, aged N = 11, n = 25; oIPSC adult N = 8, n = 13, aged N = 11, n = 17). **(e)** Optogenetically-evoked post synaptic currents for the same cells at V_h_ = -65mV (excitatory) and V_h_ = -10mV (inhibitory). Results are expressed as the mean ± SEM. (For adult group: N = 2, n = 3. For aged group: N = 5, n =7). **(f)** Representative traces of optogenetically-evoked excitatory post synaptic currents (oEPSC) after a 5ms blue light stimulation, represented as blue bar. Right: enlarged illustration of the dashed rectangle. Blue bar represents the optical stimulation. **(g)** Latency of oEPSC and oIPSC is maintained during aging for D2R-MSN in females, but oIPSC show a higher latency (around 5ms) compared to oEPSC (around 2ms). Results are expressed as the mean ± SEM. (Two-way ANOVA, Vhold: p < 0,0001, F (1, 33) = 56,88, oEPSC adult N = 3, n = 10, aged N = 7, n = 16; oIPSC adult N = 2, n = 3, aged N = 5, n = 8). **(h)** Representative traces of optogenetically-evoked excitatory post synaptic currents (oEPSC) after a 5ms blue light stimulation, represented as blue bar. Right: enlarged illustration of the dashed rectangle. Blue bar represents the optical stimulation. **(i)** Latency of oEPSC and oIPSC is preserved during aging for PV-expressing neurons in females, but oIPSC tend to show a higher latency (around 5ms) compared to oEPSC (around 2ms). Results are expressed as the mean ± SEM. (Two-way ANOVA, Vhold: p = 0,0170, F (1, 24) = 6,579, oEPSC adult N = 5, n = 13, aged N = 5, n = 10; oIPSC adult N = 2, n = 2, aged N = 2, n = 3). In all panels: significant p-values are represented in bold, N = number of animals, n = number of cells.

